# Population Genomics of *P. miniata* along the Pacific Coastline Reveal Subtly Diverging Genomics Along an Extensive Range Gap

**DOI:** 10.1101/2025.10.13.679893

**Authors:** Veronica Pagowski, Brendan Cornwell, Erik Hanson, Fiorenza Micheli, Stephen Palumbi

**Affiliations:** Hopkins Marine Station, Oceans Department, Stanford University, Pacific Grove, CA, USA; Stanford Center for Ocean Solutions, Pacific Grove, CA, USA; Department of Biology, Stanford University

**Keywords:** Bat star, *Patiria miniata*, Population genomics

## Abstract

Many marine species exhibit range gaps, patchy distributions, or genetic disjunctions throughout their ranges. These discontinuities can result from various interacting mechanisms, though directly linking ecological and historical factors to observed distributions or genetic disjunctions often proves challenging. Intriguingly, substantial research has demonstrated that these spatial and genetic discontinuities frequently occur in species with long-lived planktonic larvae, which possess the capacity for extensive oceanic dispersal. This study investigates the population genetics of one such species, the bat star *Patiria miniata*. Despite its long-lived planktonic larval stage of six to ten weeks, *P. miniata* maintains both a range gap and a, geographically separate, strong genetic disjunction throughout its distribution from Alaska to Baja California. Utilizing low-coverage whole-genome sequencing of over 200 individuals collected throughout *P. miniata*’s range between the early 2000s and 2023, we corroborate previous findings of a significant genetic disjunction across Queen Charlotte Sound, north of Vancouver Island. Additionally, we present new evidence of strong divergence at several genomic loci across an extensive range gap in Washington and Oregon, despite subtle genetic population structure at most loci here. Such differences may reflect more recent ecological or oceanographic barriers, rather than historical processes. Our results demonstrate that while marine invertebrate populations may appear panmictic based on genome-wide metrics of population structure alone, strong local selection for specific gene segments may be maintained in some populations. This research contributes to our understanding of the complex interplay between dispersal potential and local adaptation in marine ecosystems and highlights the importance of considering both genetic structure and finer-scale adaptation in coastal marine populations.

## Introduction

While the same principles of conservation biology, ecology, and evolutionary biology apply to land and marine systems, marine environments pose unique challenges that are likely to shape population genetic structure and evolution of life in the ocean (Carr et al., 2003; Kelley et al., 2016; Schiebelhut & Dawson, 2018). Larvae, for example, are a pervasive life stage at sea. While ocean currents are sometimes assumed to disperse small larvae far and wide, realized dispersal distances do not always match modelled dispersal, and frequently undershoot potential dispersal distances (Shanks, 2009).

Nonetheless, partially due to the high dispersal potential of marine larvae, past work has highlighted that marine populations tend to have higher gene flow and larger effective population sizes, decreased spatial genetic structuring, high genetic diversity, faster responses to large-scale events like environmental disturbances, and greater variability in year-to-year recruitment, compared to terrestrial species (Carr et al., 2003). In these cases, genetic drift is expected to be slow and may be outpaced by even weak selection on specific loci (De Wit & Palumbi, 2013). Despite these differences, which contribute to more open populations with high exchange rates in marine populations, genetic differentiation between populations occurs in many species, suggesting that marine populations do not always behave as large panmictic units as sometimes assumed (Sivasundar and Palumbi 2010; De Wit & Palumbi, 2013; Barney et al., 2017; Griffiths et al., 2025; Reeb & Avise, 1990; Sanford & Kelly, 2011; Palumbi et al. 2023; St. John et al., 2025; Van Wyngaarden et al., 2017).

A surprising finding of a large collection of work from recent decades suggests that genetic disjunctions occur in many marine species, even in organisms with long-lived planktonic larvae, and sometimes in the absence of obvious dispersal barriers (Sivasundar and Palumbi 2010; El Ayari et al., 2017; Gardner & Wei, 2015; M. Liu et al., 2012; Reeb & Avise, 1990; Shahdadi et al., 2022; Toms et al., 2014). These genetic breaks can occur in different locations, depending on the species (Kelly & Palumbi, 2010), highlighting species-specific responses to mechanisms that contribute to genetic disjunctions. Marine species’ distributions can likewise be patchy, but the locations where range gaps and genetic breaks occur are not always the same (Brasher et al., 1992; Gaffney et al., 2007; Herranz & Leander, 2016; Hunter & Halanych, 2008; Kirkendale et al., 2004; Metz et al., 1998; Vecchioni et al., 2019), highlighting the complexity of historical and ecological factors that underlie genetic structuring and patchy ranges in marine species (Pagowski & Micheli 2024).

Effective stock conservation and management of marine systems can be improved by incorporating the population genetic dynamics and estimates of connectivity, which has been highlighted as key for designing effective conservation and restoration strategies in marine ecosystems (Balbar & Metaxas, 2019; Carr et al., 2017; Christie et al., 2010; Lagabrielle et al., 2014; Schill et al., 2015; Sullivan-Stack et al., 2022; Wright et al., 2015; Haupt et al. 2013, Munguia-Vega et al. 2015). Thus, additional studies documenting population genetics and genomics of marine species across broad geographic ranges are needed. As large-scale sequencing is now becoming more financially feasible in non-model organisms, such studies are just beginning to inform conservation strategies (Jeffery et al., 2022). Here, we examine the population genomics of the bat star *Patiria miniata*, a species that encompasses a genetic disjunction, an extensive range gap, and a strong oceanographic boundary within its range.

The bat star *Patiria miniata* has a long-lived planktonic larval stage, with larvae remaining in the water column for six to ten weeks before metamorphosing into benthic juveniles (Basch, 1996; Rumrill, 1989; Strathman, 1987; Sunday et al., 2014). Its range of distribution spans from Sitka, Alaska to the Revillagigedo Islands in Mexico (Kozloff, 1983; Lambert, 2000), though it is rarely found south of Northern Baja California (iNaturalist community, 2025). Despite this extended larval duration, this species shows a strong genetic disjunction in the northern part of its overall range (Keever et al., 2009), as well as an extensive range gap across Washington State and Oregon (>500km). The species also is abundant across a significant oceanographic boundary in southern California at Point Conception, also known by its original Chumash name *Humqaq*. Thus, for a broadly distributed, abundant species with a long larval duration, *P. miniata* shows surprising discontinuities, both spatial and genetic.

These features make this species an exciting model to investigate how different forces shape the population distribution and genetics of marine invertebrates with long-lived planktonic larvae. Additionally, the availability of a limited set of historical samples in this study enables us to address questions about the temporal variability of genomics in this species before and after the sea star wasting disease event along the west coast of North America. Sea star wasting disease devastated over twenty species of sea stars but left *P. miniata* populations comparatively unaffected (Dawson et al., 2023; Hewson et al., 2018; Menge et al., 2016; Miner et al., 2018; Montecino-Latorre et al., 2016). However, the potential effects on genetic structure and diversity of *P. miniata* have not been addressed.

In this study, we employ low-coverage whole-genome sequencing (lcWGS) to investigate the demographic history, population genetic structure, and selection dynamics in *P. miniata* and explore these results in the context of demographic history and oceanography. Specifically, we asked (1) whether *P. miniata* populations from Haida Gwaii to Baja California are genetically variable, (2) if genomic differentiation occurs across an extensive range gap in Washington and Oregon, and (3) if specific genomic regions are divergent between samples collected at the same location in different years.

## Methods

### Population Sampling and DNA Extraction

We sampled *P. miniata* populations throughout much of center of their reported geographic range, which spans from Sitka, Alaska to the Revillagigedo Islands in Mexico (Kozloff, 1983; Lambert, 2000). In 2023-2024, we collected samples from Ucluelet (southern Vancouver Island) to northern Baja California (**Supp. Table 1**). **Figure 1** summarizes the locations and collection source for all populations included in this analysis. At each sampling location, we collected at least 12 samples for lcWGS sequencing. In the field, 2-8 tube feet from live individual specimens of *P. miniata* were collected at low tide and placed into tubes with 90% or greater ethanol. In the southern part of the range (San Diego, Santa Barbara, Baja California) live specimens were rare in the intertidal and tissue samples collected in this study were obtained by sampling specimens found by SCUBA diving. Additionally, aquarium tissue samples were obtained similarly from specimens originally collected in Fort Bragg, Santa Barbara, Los Angeles, and San Diego (kindly donated from the Noyo Center for Marine Science, Santa Barbara Sea Center, Heal the Bay Aquarium in Los Angeles, and Birch Aquarium in San Diego). These *aquarium* samples also included tube feet collected from live specimens, that were subsequently stored in ethanol. While these samples were collected in a range of years, our best approximation is between 2016-2017 in Fort Bragg, 2008 in Santa Barbara, and 2016 in San Diego. Samples from Los Angeles were collected at an unknown date, likely 10+ years ago. While exact dates of collection are not known for these samples, they provide valuable context into the potential year-to-year variability of population genomics of bat stars along their range when compared to samples we collected in 2023-2024. *Historical* samples were kindly donated by the Hart laboratory at Simon Fraser University and resequenced in this study. These samples include bat star tissue from several northern locations (Haida Gwaii, Central BC, Winter Harbor, and Bamfield) as well as a small number of samples still available from southern locations (Fort Bragg, San Diego). These *historical* samples were stored in a -20 degree freezer since the early 2000s. Throughout this manuscript *new* samples refer to those collected in 2023, *historical* samples refer to collections from the early 2000s and *aquarium* samples refer to sample tissue donated by several aquaria. In addition, sex of sampled individuals was available for a small subset of 10 samples collected in Monterey. DNA extraction for all samples was performed according to manufacturer instructions using an Omega Bio-Tek E.Z.N.A. Tissue DNA Kit.

**Figure 1:**
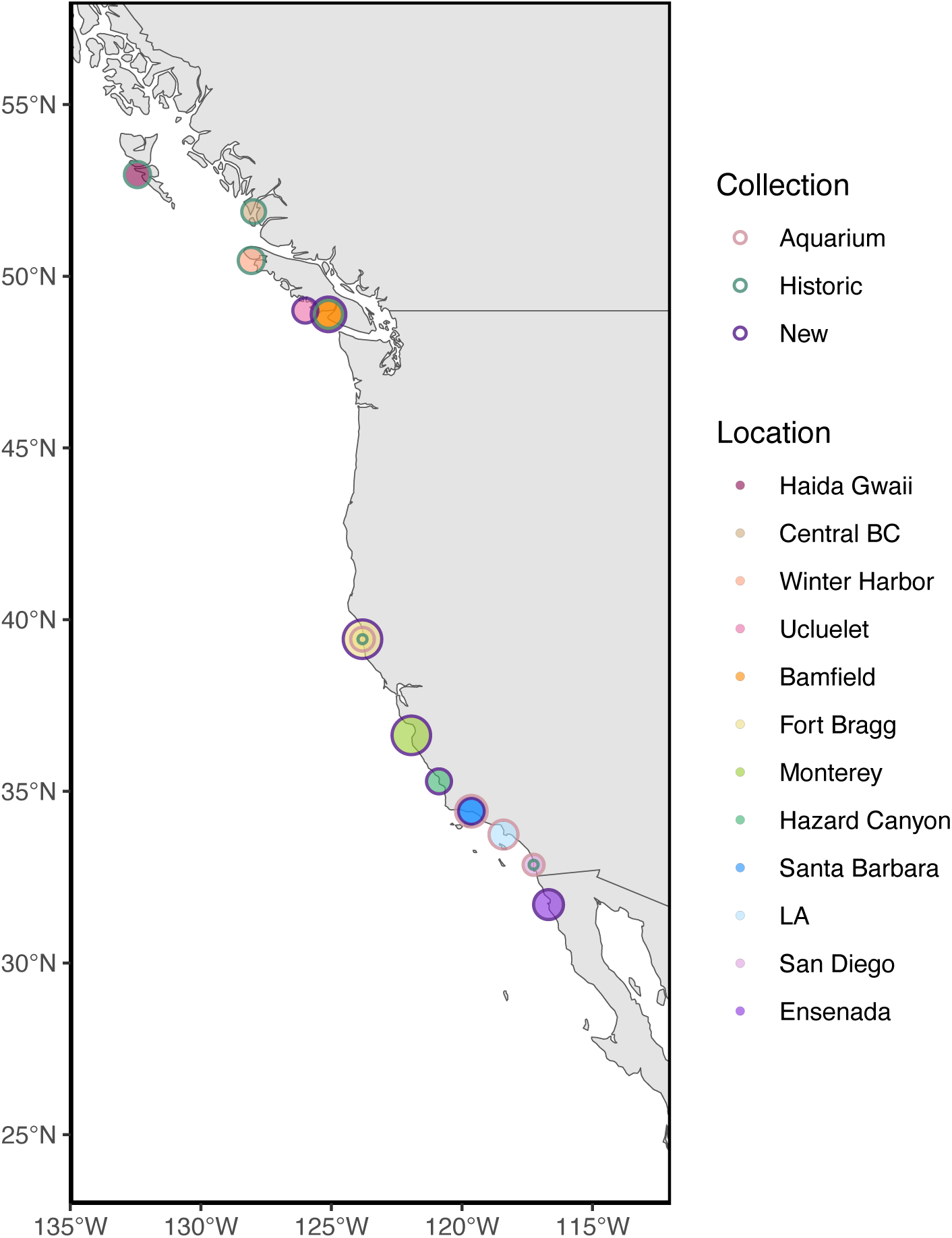
Sampling Locations. Points represent a sampled populations and point outline color corresponds to the collection type (*new*, *historic*, or *aquarium* samples). Points with multiple outline colors represent locations where multiple sample types were available. Sample size is represented by point size (4-28 samples per population). Except for *historic* Fort Bragg and San Diego locations, all sample sizes are greater than 8. A table of sampling locations and sample sizes is also available in. **Supp. Table 1.**

### Sequencing

Following DNA extraction, sample library preparation and sequencing were performed by Texas A&M AgriLife Genomics and Bioinformatics Service on two lanes using a NovaSeq X 10B (300 cycles/lane). Scripts for all filtering and analysis steps are available [here](https://github.com/vpagowski/Pminiata_popgen_paper.git). In total, we sequenced 248 *P. miniata* samples at low coverage (avg. of 5.13x), as well as 9 individuals at high coverage (>18x). All samples were sequenced in one of two total sequencing rounds, with 8 re-sequenced individuals included in the second sequencing batch to estimate lanes effects. While most of our analyses utilize data from hard calls generated with GATK and further analyzed with other software, we also utilize imputed genotype data generated with ANGSD in some analyses (see below for details on methods for both approaches). In most cases, if imputed or hard call results are shown in the main figures, an analogous analysis is included in the supplementary materials.

### Read Filtering and SNP Calling

Prior to calling SNPs, we used Skewer (Jiang et al., 2014) to trim adapters and filter reads for quality using a quality cutoff of 20, end quality of 28, and a custom set of adapters to remove common adapters and several highly repetitive sequences that were very abundant. We verified that sequences were adapter-free with FastQC (Andrews et al., 2010). Following this basic quality filtering, Burrows-Wheeler Aligner (BWA) (Li & Durbin, 2010) was used to align reads to the [P. miniata genome](https://www.ncbi.nlm.nih.gov/refseq/annotation_euk/Patiria_miniata/100/). SAMtools (Danecek et al., 2021) and Picard (*Picard Tools - By Broad Institute*) were used to sort reads, remove duplicates, call read groups, index reads, and assess alignment coverage.

To call variants, we applied GATK best-practice protocols for variant discovery and joint genotyping (Auwera & O’Connor, 2020; DePristo et al., 2011; McKenna et al., 2010). For subsequent filtering and linkage disequilibrium pruning we used PLINK (Purcell et al., 2007) and VCFtools/BCFtools (Danecek et al., 2011). We removed indels, retained biallelic sites only, and tested a series of filtering steps to ensure the stability of our results. In our main figures we use the following filters: maximum per-site missingness rate of 50% for each population, max sequencing depth of 100x, minimum quality of 20, minor allele frequency of 0.05. We pruned SNPs in linkage disequilibrium using 200kb windows in steps of 20kb with r^2^ values of 0.5 or higher. Using our GATK pipeline, we retained 8,338,770 hard-call SNPs after all filtering steps. Using genotype imputation, we retained 30,008,952 SNPs, using a similar set of filters. **Supp. Table 2** contains additional information on filtering and SNPs retained for each analysis.

We also used ANGSD (Korneliussen et al., 2014) to generate genotype likelihoods using the GATK model to infer major and minor alleles from genotype likelihoods (-doMajorMinor 1) and assume fixed major and minor alleles (-doMaf 1). To generate our NMDS plot with imputed genotypes, we also pruned variants in linkage disequilibrium using ngsLD (Fox et al., 2019), using previously described protocols (Lou et al., 2021). To this end, we generated an initial set of independent sites by pruning SNPs with an r^2^ of 0.6 or greater at a maximum distance of 50kb or 50 SNPs (--max_kb_dist 50, --max_snp_dist 50, --min_weight 0.6). Using this set of independent sites, we regenerated our NMDS based on genotype likelihoods, using allele frequency as a prior and filtering for SNPs in a minimum of 117 individuals (50% of our dataset), minimum minor allele frequency of 0.05, minimum mapping quality of 20, minimum base quality of 20, and SNP p value of 1e-6 of lower.

To investigate sex differences using imputed genotypes, we generated an equivalent set of SNPs using all the same settings but filtered for a minimum of 10 individuals present for each SNP instead of 50% of the dataset. This generated 38,042,207 SNPs and very similar overall results. Using hard call genotypes, we document a slight batch effect between sequencing on two lanes after investigating 8 resequenced individuals. This effect is much less severe using imputed genotypes. Thus, throughout this manuscript we use both approaches to inform our results, replying mostly in imputed genotypes for overall visualizations of population structure such as NMDS and admixture plots.

### Admixture

We conducted admixture analysis with genotype likelihoods using ngsADMIX (Skotte et al., 2013), using previously filtered and linkage-disequilibrium pruned genotypes. We used 500 maximum iterations for K=2-4 groups in ngsADMIX.

### F_ST_ and Population Diversity Metrics

We used PLINK to calculate pairwise F_ST_ between populations with sample sizes greater than 8 (all populations except a small number of *historical* samples from Fort Bragg and San Diego), using the Weir and Cockerham method (Weir & Cockerham, 1984). Per-population Tajima’s D, and nucleotide diversity metrics were similarly calculated with PLINK in 10kb windows. Per-sample inbreeding coefficients were also calculated in PLINK.

### Outlier Analysis and GWAS

To investigate if specific genomic regions were highly differentiated across Queen Charlotte Sound, north and south of Point Conception/Humqaq, and north and south of *P. miniata’s* expansive range gap, we plotted F_ST_ generated in PLINK, using individuals pooled across these different regions where we hypothesized local genomic differentiation could occur. We also conducted an analysis in SNPEff (Cingolani et al., 2012), using the *P. miniata* genome, to identify predictions about gene impact. Plots were made in ggplot2 (Villanueva & Chen, 2019) with R version 4.2.3 (R Core Team, 2021). We then used PLINK to run a genome-wide association study (GWAS), with latitude of collection as a response variable. Covariates were included in the analysis by generating an additive linear model with PCs 1 and 2 from our hard-call PCA **(Supp. Figure 1)** and sequencing batch as explanatory variables for sample latitude. Residuals for this model were used as a response variable rather than latitude in our covariate-corrected GWAS.

### Signatures of Selective Sweeps

To look for evidence of selective sweeps, we used PLINK to generate single scaffold VCF files and then used VCFtools to output raw allele count data. We then used Sweepfinder2 (DeGiorgio et al., 2016) to generate sweep likelihoods for scaffolds in which we observed strong signatures of differentiation from earlier GWAS and outlier analysis results. BEDtools (Quinlan, 2014) and custom scripts using BLAST (Sayers et al., 2021) were used to identify annotated genes within our strongest selective sweep peaks and BLAST hits with an e value 0.001 or less and maximum best targets of 8 hits. Plots of these results were made in R using ggplot2.

### Demographic History

Demographic history was inferred using PSMC (Li & Durbin, 2011) and Beta-PSMC (J. Liu et al., 2022). While both methods rely on the same principle of inferring ancestry based on heterozygote densities throughout the genome (Li & Durbin, 2011), Beta-PSMC provides more detailed information on more recent timescales (J. Liu et al., 2022). For PSMC, we tested several time interval parameters to test for consistency of our results, but use a simulation run with 25 iterations (-N), a maximum coalescent time of 15 (-t), and the interval pattern parameter of 26*2+4+7+1 in main figures. We generated 100 bootstrap replicates by splitting and subsampling the fasta alignment generated with the split utility within PSMC. For Beta-PSMC, we use the default interval pattern parameter of 20*1, initial theta/rho of 5, maximum coalescent time of 15, and 25 iterations, similarly generating 40 bootstrap replicates. Since one Haida Gwaii sample sequenced at significantly higher (∼50x) coverage (**Supp. Figure 2**), compared to our other samples, we randomly down-sampled our bam alignment file for this individual to more closely match the coverage of our other samples (35x), as results of demographic analyses are sensitive to genome coverage (**Supp. Figure 2,3**). Results are scaled with a mutation rate of 9.13e^-9^. This value aligns with the estimated germline mutation rate for the crown-of-thorns sea star (Popovic et al., 2024). Notably, this value is higher than previously estimated mitochondrial mutation rate of 3.8e^-9^ in *P. miniata* (McGovern et al., 2010). The true germline mutation rate and generation time are not well known for *P. miniata*, so we caution that the precise time period of the divergence between populations might differ. We highlight that changes to these parameters are likely to shift the timeline of divergence, but not the overall shape of the PSMC curves (S. Liu & Hansen, 2017).

### Supplemental Analyses

In addition to data presented in the main figures, we conducted a series of additional analyses presented in the supplemental data. To further investigate how specific genomic regions may drive differentiation between our populations, we conducted a windowed PCA, utilizing the WinPCA software (Blumer et al., 2025). This allowed us to visualize how specific genomic regions contribute to driving variation among samples. To investigate how oceanography and long-term dispersal may contribute to the genetic patterns observed, we utilized Opendrift, an open-source Python-based framework for Lagrangian particle modelling (Dagestad et al., 2018), to model larval dispersal over a 10-week period in several El Nino (2016,2020,2023) and La Nina (2000, 2010, 2011) years, using best approximations for parameters describing larval swimming and growth over time. We also investigated signatures of past migration and directionality between populations using Treemix (Pickrell & Pritchard, 2012), after generating a consensus tree with Phylip (Felsenstein, 1993) to describe relationships among populations. More details on each of these analyses can be found in the **Supplemental Methods** document.

## Results

### Population Genetic Structure

Our analyses first examined whether genetic breaks or strong population structure are evident along the geographic range of *P. miniata.* To this end, we visualized broad patterns in our low-coverage samples with non-metric multidimensional scaling (NMDS) and principal component analysis (PCA).

Corroborating past work on this species (Keever et al., 2009), we document a very strong genetic split between populations north of Vancouver Island. These samples include *historical* samples from Central BC and Haida Gwaii. Both of these populations are highly divergent from any populations south of Queen Charlotte Sound. We also find that while *historical* and *new* samples from Bamfield generally cluster together in NMDS space, a handful of *new* Bamfield samples show high differentiation from all other samples (**Figure 2A**). Interestingly, these divergent samples come from both of our nearby sampling locations at Bamfield. To investigate whether population structure could be detected if these strongly divergent populations and individuals were removed, we regenerated the NMDS plot using a subset covariance matrix generated from ANGSD that excluded Haida Gwaii, Central BC, and outlier Bamfield samples.

**Figure 2:**
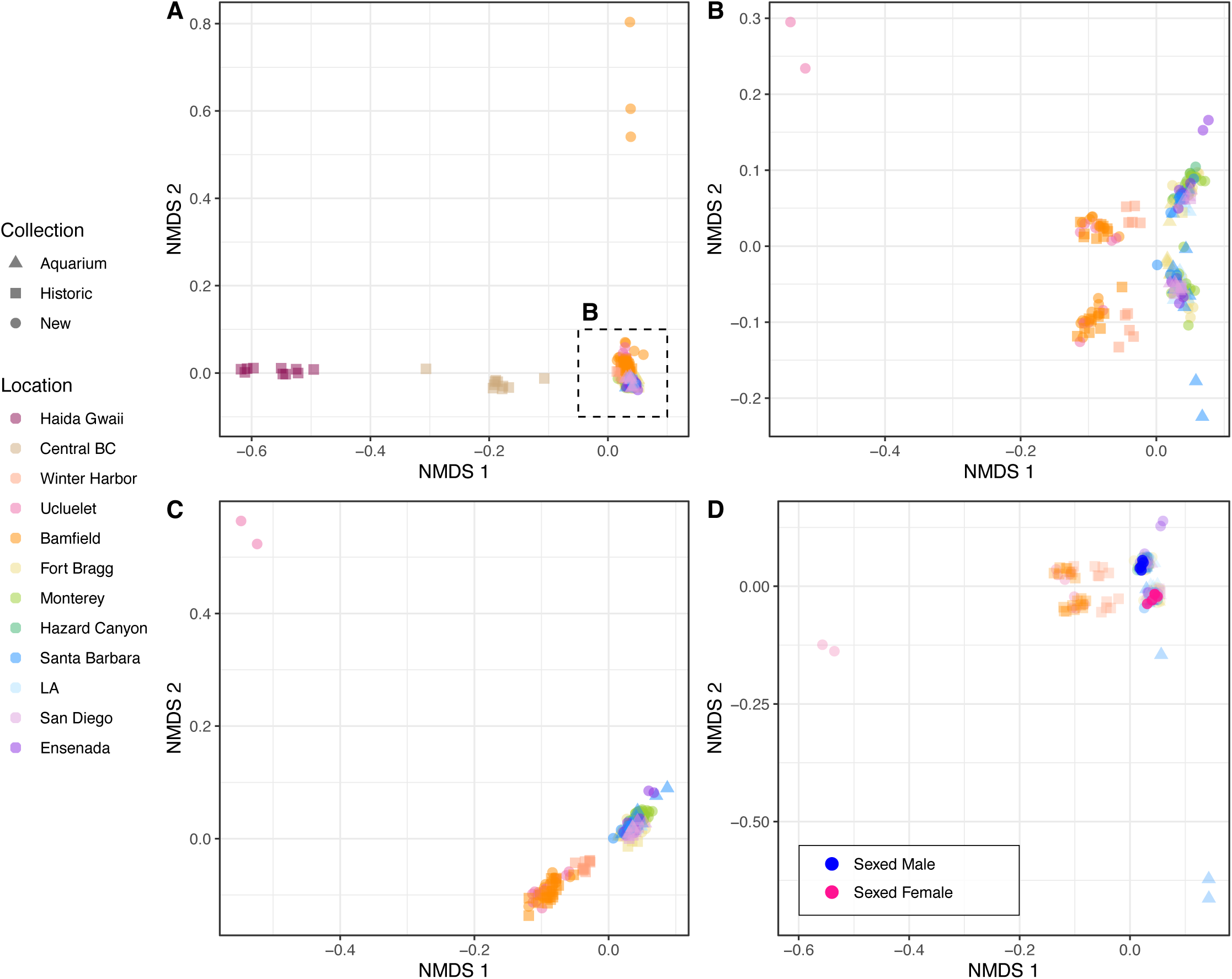
NMDS Analysis of Imputed Genotypes. NMDS plot showing *aquarium*, *historic*, and *new* samples (triangles, squares, and circles, respectively) highlighting a strong genetic disjunction across Queen Charlotte Sound **(A)**. Re-calculating NMDS for a subset of samples (dashed square in A) results in separations of Vancouver Island samples from California and Baja California samples along PC1 and an approximately even subset of samples from all populations separated along PC2 **(B)**. This separation is no longer visible when two scaffolds that differed between sexed (64 and 65) are removed from the analysis **(C)**. When re-running our analysis with 10 individuals of known sex, this time using a minimum individual filter of 10 rather than 50% of the data, similar results are obtained and sexed males and females map of opposing clusters along NMDS 2 **(D)**.

Using this approach, we detect a distinct genetic signature of the range gap, such that samples from California form a cluster distinct from samples collected in Vancouver Island at Bamfield, Ucluelet, or Winter Harbor (**Figure 2B**). However, using this analysis, we find that samples from Winter Harbor were more similar to southern samples even though they are geographically closer to Haida Gwaii and Central BC.

Within the southern clusters there are multiple outlier individuals. Namely, 2 *new* individuals from Ucluelet, 2 *new* samples from Ensenada, and 3 *aquarium* samples from Santa Barbara. Population structure signals appear to be so weak south of the range gap that high similarity of closely related individuals might drive this, as removal of select outlier individuals often results in new outliers appearing.

### Possible sex-specific genetic signals

NDMS plots (**Figure 2B**) split geographic samples along axis 2 between all locations: because these splits have almost equal sample sizes, we hypothesized that sex-differences might contribute to this divergence. To address this hypothesis, 5 sexed males and 5 sexed females were sequenced in our second round of sequencing (which also included 56 other samples, including resequenced individuals). We found that males and females were most divergent at two scaffolds **(Supp. Figure 4)**. When these two scaffolds were removed, the split across NMDS 2 was not observed (**Figure 2C**). However, when included, known males and females mapped to opposing clusters (**Figure 2D**).

While a mechanistic explanation of these differences is beyond the scope of this work, we find that one of the scaffolds at which putative females and males differ is particularly heterozygous (**Supp. Figure 4,5**). Furthermore, putative females share some regions of homozygosity that are not found in putative males within this divergent scaffold, suggesting that this scaffold could correspond to an unassembled sex chromosome. An analysis of linkage disequilibrium at different scaffolds supports this hypothesis as highly linked loci appear throughout this scaffold (**Supp. Figure 6**). As all of our scaffolds have similar coverage across samples and the reference genome for this species comes from a sperm sample, this could be suggestive of a ZW system, in which females are characterized by two differing chromosomes and males possess two copies of the same chromosome. However, a more complex system is also possible, as we do see other regions that are differentiated throughout the genome between males and females, though not as strongly. Additionally, we do not observe that our putative males have 2x the coverage of putative females at this scaffold as would be expected with the above hypothesis. Putative male coverage at this scaffold was only marginally higher than putative female coverage (4.49x and 4.21x respectively for males and females). Furthermore, heterozygosity was comparable for both males and females **(Supp. Figure 5)**.

### Northern affinity of southern outliers

Using hard call data to identify population structure using principal components analysis, we obtain generally similar results. However, we also find that *aquarium* samples from both Fort Bragg and Santa Barbara collected between 2008 and 2017 sometimes form a unique cluster that overlaps with some of our *new* northern samples from Bamfield and Ucluelet and *historical* and *aquarium* samples from Fort Bragg (**Supp. Figure 1**). While speculative, this could suggest that southern samples may have historically received more northern input. Our windowed PCA analysis provides additional support for this hypothesis, as we find that at one of our most differentiated genomic regions (SC 72) across the range gap, all of our *aquarium* samples from Fort Bragg (and a couple *aquarium* samples from Santa Barbara) show an intermediate signal of differentiation (**Supp. Figure 5E**). In this analysis we also sequenced a small number of *historical* individuals from Fort Bragg (n=4) and San Diego (n=4). As only a small number of individuals were available from these populations from an earlier study, we exclude these samples from our analyses involving population comparisons. However, we note that *historical* samples from Fort Bragg cluster most closely with our *aquarium* samples from Fort Bragg and Santa Barbara, also collected in the past while *historical* San Diego samples cluster with other southern counterparts, further supporting the idea of subtle shifts in population structure over time.

### Admixture

Admixture analyses with ngsAdmix highlight the strong genetic disjunction across Queen Charlotte sound and the subtle genetic divergence across the range gap. Using k=2 groups only shows the disjunction across Queen Charlotte Sound, with Central BC samples having ancestry resembling both groups. Using k>2 highlights the genetic signature of differentiation across the range gap between Bamfield and Fort Bragg (**Figure 3**). Notably, several Bamfield samples show unique genetic signature signatures in this analysis, as was also observed in our NMDS plots and PCA. Broadly, we highlight that this analysis further corroborates the strong disjunction across Queen Charlotte Sound and weak, but detectable, genetic divergence across the range gap.

**Figure 3:**
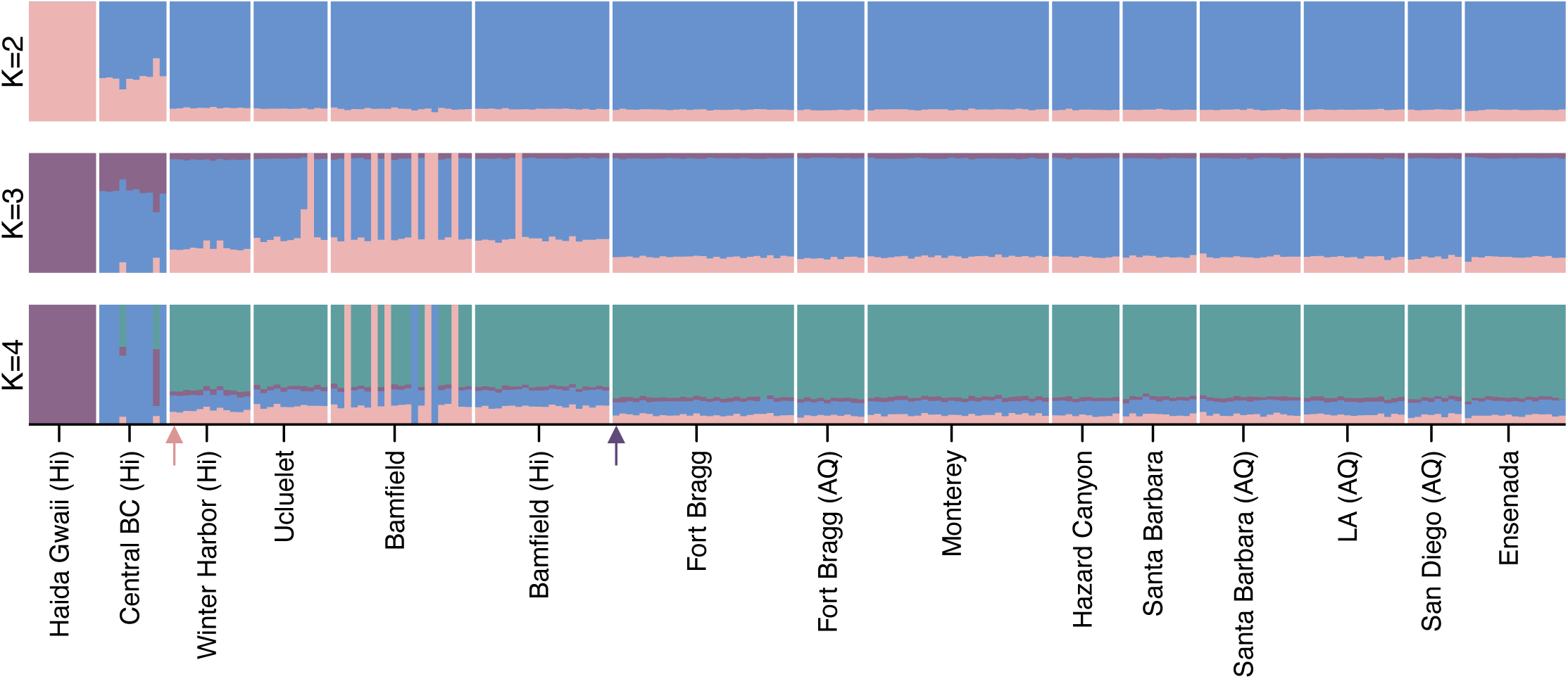
Admixture Analysis. All data in this figure represent analysis of imputed genotypes for populations with a sample size of 8 or more. Results represent admixture analysis using k groups of 2-4. Sample labels with (Hi) or (AQ) indicate historic and aquarium samples, respectively. Sampling locations are arranged from north to south (left to right). Pink and blue arrows correspond to the genetic disjunction and range gap, respectively.

### Population Differentiation and Genetic Diversity Metrics

To quantify genetic differences at each SNP between each population and measure population wide nucleotide diversity, we calculated pairwise F_ST_ by geographic site as well as several metrics that provide insight into genetic diversity (nucleotide diversity, Tajima’s D, inbreeding coefficient). To this end, we used our hard call dataset to calculate F_ST_ between all populations with a sample size greater than 8 individuals. Again, we find a strong genetic split between populations from Haida Gwaii and Central BC and subtle genetic differentiation across the range gap (**Figure 4A**). South of the range gap, populations are effectively indistinguishable from each other using F_ST_ metrics, with F_ST_ values between these populations range from 0 to 0.006 (**Figure 4A**).

**Figure 4:**
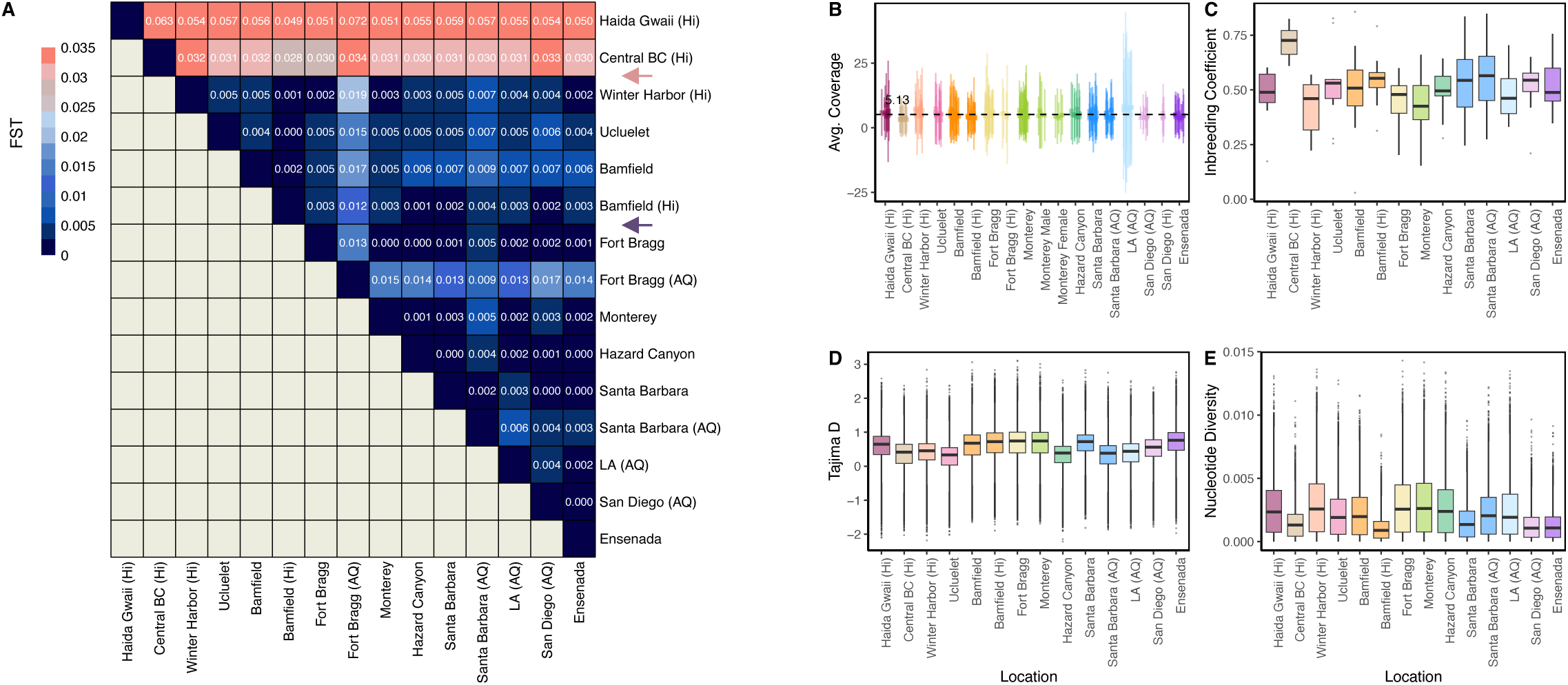
F_ST_, Sequencing Coverage, and Diversity Metrics of Populations. All data in this figure represent analysis of hard calls (rather than imputed genotypes) for populations with a sample size of 8 or more. The matrix represents pairwise F_ST_ between all populations, with (Hi) indicating historic samples and (AQ) representing aquarium samples collected prior to 2018. All other samples were collected in 2023 **(A)**. Average per-sample coverage and standard deviation **(B)**, inbreeding coefficients **(C)**, Tajima’s D calculated in 10kb windows, **(D)** and Nucleotide diversity calculated in 10kb windows is shown in the right panels. In each box and whisker plot, the median is indicated by a solid line and outliers are indicated by points. *D* and *E* represent pooled population averages from locations along the genome while *B* and *C* represent single values per individual. The dashed line and label in *A* represent the average coverage of 5.13x and the pink and blue arrows in A represent the locations of the genetic disjunction and range gap, respectively. From left to right, locations are north to south, with duplicate colors representing a different collection type (Hi/AQ) for the same location.

Notably, we also find that when *aquarium* samples from Santa Barbara and Fort Bragg are compared to other populations, F_ST_ values are higher (up to 0.009) than when *new* samples from these populations are compared to other populations (up to 0.007). However, as some of these *aquarium* samples from Fort Bragg had higher coverage variation and much lower genetic diversity indices compared to most of our other samples (**Figure 4B**), these are removed from main figures to avoid overinterpretation of results related to these samples. Analogous figures including these samples are available in **Supp. Figure 7**.

Overall, we find that nucleotide diversity and Tajima’s D values are similar throughout most of the species’ range (**Figure 4D,E**). However, we note a low nucleotide diversity and high inbreeding coefficient for samples collected from Central BC (**Figure 4C,E**).

### Outlier Analysis and Evidence of Local Selection

To investigate if specific genomic differences might drive the patterns of genomic differentiation that we observed above, we conducted an outlier analysis and quantified per SNP F_ST_ along the genome. When comparing populations north and south of Queen Charlotte Sound, we find that high overall F_ST_ (0.021), calculated as the weighted average F_ST_ between Vancouver Island populations and those north of Vancouver Island, is distributed throughout the genome (**Figure 5A**), suggesting that genetic drift may have accumulated after long-term divergence of these populations. Note in figure 5 that the available *Patiria miniata* genome has been assembled as scaffolds only, that have not yet been assembled into chromosomal lengths.

**Figure 5:**
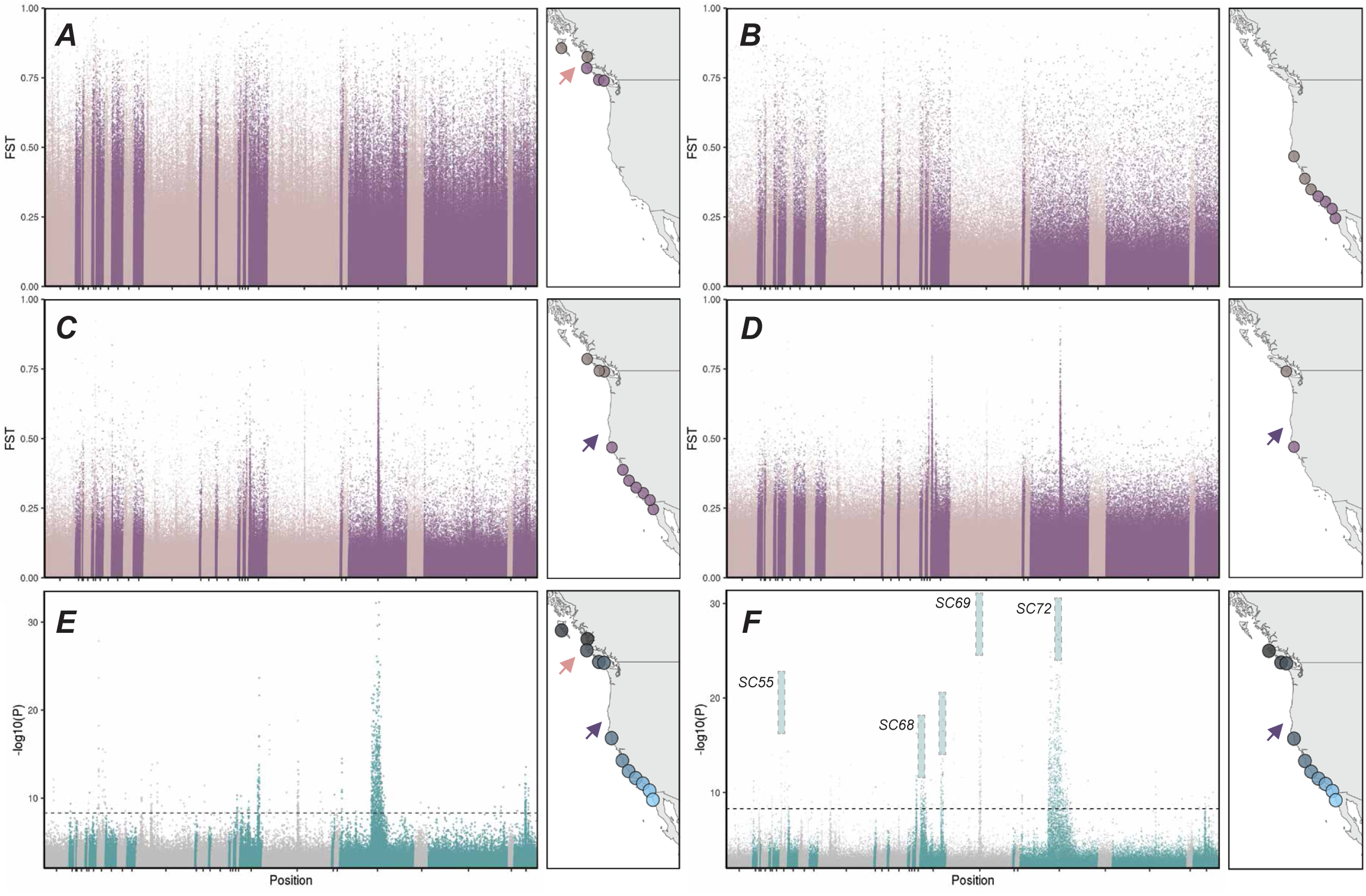
Outlier Analysis and GWAS Results. All data in this figure represent analysis of hard calls (rather than imputed genotypes) for populations with a sample size of 8 or more. The top panel represents calculated F_ST_ along the genome, comparing populations north and south of Queen Charlotte Sound **(A)**, and north and south of Point Conception (excluding samples from Canada) **(B)**. The middle panel shows F_ST_ differentiation north and south of the range gap, including all populations south of Queen Charlotte Sound **(C)** or only *new* Bamfield and *new* Fort Bragg samples **(D)**. The bottom panel shows GWAS results using sampling site latitude as a response variable using all populations and no covariates **(E)** or all populations excluding Haida Gwaii and Central BC, using PCs 1 and 2 and sequencing round as covariates **(F)**. Dashed lines indicate Bonferroni-corrected significant threshold. Color changes indicate different scaffolds and F_ST_ values below zero are not shown. The maps to the right of each plot indicate the populations being analyzed in each figure. Labels in *F* indicate scaffold ids for genomic regions with strong differentiation in the GWAS. The pink and blue arrows represent the locations of the genetic disjunction and range gap, respectively.

When we compare populations across Point Conception/Humqaq (only including individuals collected south of the range gap), we again find that there are no clear signals of differentiation at specific genomic loci (**Figure 5B**), suggesting that this oceanographic boundary might not be highly relevant for *P. miniata*. By contrast, when populations north and south of the range gap are compared (excluding our highly divergent Central BC and Haida Gwaii samples), we find that signatures of differentiation are localized to specific genomic regions (**Figure 5C**). In particular, differentiation at scaffold 72 stands out in comparing across the range gap.

The strongest signatures of these localized genomic differences can be observed when comparing the two *new* populations flanking the range gap, Bamfield (n=22 ) and Fort Bragg (n=28) (**Figure 5D**). In addition, these signals of local differentiation are highly evident in a genome wide association analysis, using collection latitude as a response variable (**Figure 5E,F**). When all samples are compared, SNPs localized to several genomic regions are strongly associated with latitude including differentiation at scaffold 72 (**Figure 5E**). After removing divergent Haida Gwaii and Central BC samples and correcting for covariates (sequencing batch and first 2 PCs), we continue to detect multiple regions where genomic differentiation is strongly associated with latitude (**Figure 5F**). These regions include scaffold 72 as well as scaffolds 55, 68, and 69. Because these signals disappear if samples north of the range gap are excluded (**Supp. Figure 8)**, we conclude that this differentiation is dominated by genomic differences north and south of the range disjunction.

### Evidence of Selective Sweeps

One of the most striking findings in this study is the strong signatures of localized genomic differentiation in scaffold 72 between populations north and south of the range gap, despite fairly low overall population structure across the rest of the genome. To investigate this region and others, we used Sweepfinder2 (DeGiorgio et al., 2016) to look for evidence of selective sweeps within populations. We find strong evidence of a selective sweep within scaffold 72 in almost all (all except Central BC) populations north of Fort Bragg (**Figure 6**) and no evidence of sweeps within populations south of the range disjunction in this scaffold. These geographic differences suggest that populations north of the range gap have experienced local adaptation and directional selection, and that these genetic changes are concentrated in scaffold 72. This region was also associated with high linkage disequilibrium and excess low frequency variants, characteristics of a strong selective sweep (**Supp. Figure 9**). This region also aligns with high F_ST_ between populations north and south of the range gap.

**Figure 6:**
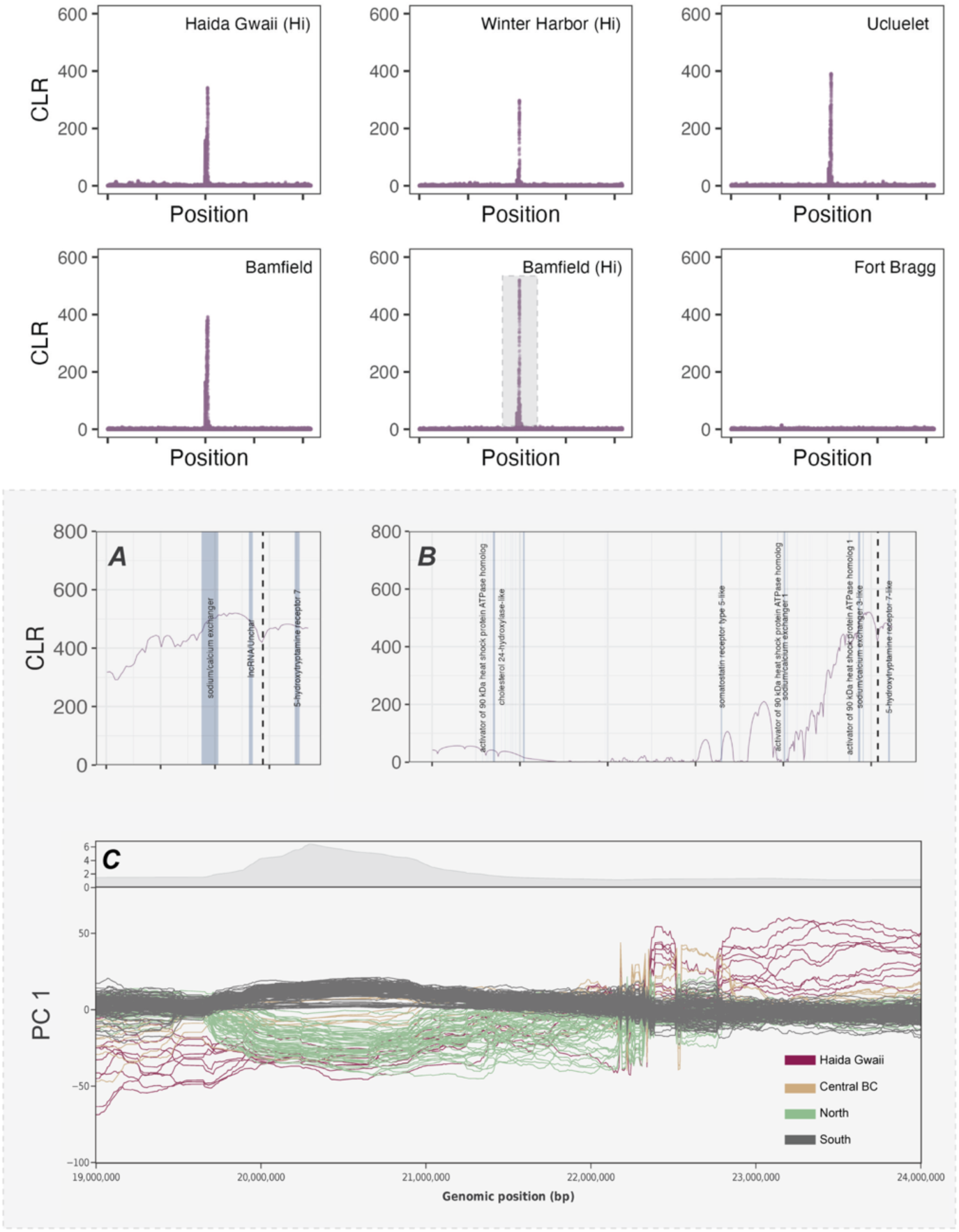
Evidence of Selective Sweeps. We detected strong signatures of selective sweeps in many populations and a more extensive analysis of these is available in **Supp.** Figure 10. This figure highlights a particularly strong sweep identified in most (all but Central BC) populations north of the range gap, both *new* and *historic* using Sweepfinder2. High cumulative likelihood ratios (y axis) indicate strong evidence for selective sweeps within populations. The top panel shows SweepFinder2 results for populations north (Haida Gwaii, Winter Harbor, Ucluelet, Bamfield) and south of the range gap (Fort Bragg and Monterey). The sweep is also detected in *historic* Bamfield samples and never detected in any population south of the range gap. The bottom panel shows a zoom in of the region highlighted in the top panel, with genes annotated in the genome intersecting with the tallest portion on the peak **(A)**. A more extensive region with annotations from blast searches for top hits intersecting with the highest sweep likelihoods is shown to the right **(B)**. For hits that have multiple mRNA matches (i.e. multiple exons), a gene label only appears for the first match. The dashed line in *B* indicates the SNP location that is most common in Bamfield samples and most rare in Fort Bragg samples (the two populations flanking the range gap) in a site-frequency analysis. Scale bars indicate 10 Mb intervals in top panels, 20 kb in panel A, and 100kb in panel. Results from a windowed PCA analysis of the highlighted region are shown in **(C)**. Y axis represents PC1, and the percentage of variance explained by PC1 along the region of interest is highlighted in the top grey plot. Plot C was generated with WinPCA software.

While scaffold 72 region is fairly large (spanning over 500kb) and encompasses multiple genes, we highlight that the strongest signatures of the selective sweep intersect with three genes annotated in the *P. miniata* genome (**Figure 6A**) including a putative heat shock protein activator, a sodium/calcium exchanger, and a serotonin receptor. BLAST results for a more expansive region surrounding this sweep also include genes associated with cholesterol/hormone production/and cell signaling (**Figure 6B**). When we investigated this region with a windowed PCA analysis (**Figure 6C, Supp. Figure 5E**), we found that samples north and south of the range gap are divergent at this genomic location.

We find similar signatures of selective sweeps at different loci in multiple populations. For example, a sweep in the distal half of scaffold 68 is visible in 5 of 9 populations south of the range gap (**Supp. Figure 10**). Strong signatures of selective sweeps are also evident at several additional loci in populations north of the range gap such as a sweep in scaffold 66 in Haida Gwaii, Bamfield, and Ucluelet, (**Supp. Figure 10**).

### Temporal Patterns in Selection

In this study, we wanted to investigate whether genomic signatures of the sea star wasting disease event could be detectable by comparing samples collected across multiple time points. Broadly, we did not observe strong evidence of genetic differences that could be attributed to selection between time points. For example, when comparing our *historical* or *aquarium* samples with *new* samples from the same location, we do not observe strong genetic differentiation at specific loci, suggesting that strong localized selection after wasting disease either did not occur or was not detectable in this analysis. Rather, differences between *historical* and *new* samples appear to be distributed throughout the genome (**Supp. Figure 11**).

### Impact of Polymorphisms

To further investigate differentiated regions between populations, we conducted an analysis in snpEff to identify the impact of SNPs in several of our most highly diverged scaffolds (**Supp. Table 3**). This analysis highlights that, while a few SNPs may have a moderate impact on gene function, many of the observed differences between populations occur in intergenic regions. We also interrogated SNPs with high F_ST_ (comparing populations across the range gap excluding our highly diverged Haida Gwaii and Central BC samples) throughout the genome, by identifying outliers in a F_ST_ vs heterozygosity plot (**Supp. Figure 12**) and identifying GO terms for our outlying variants. This analysis highlights differences in genes associated with ion and small molecule binding and cation transport.

### Demographic History

We next investigated how historic changes to population sizes might inform our results of genetic differences across Queen Charlotte Sound and across the range gap. We conducted demographic modelling to infer individual ancestries and investigate how historical changes to populations may have contributed to existing patterns. Results suggest a population bottleneck in our two samples north of Vancouver Island at the same time that populations south of Queen Charlotte Sound, including populations north of the range gap, experienced population expansions.

PSMC analysis was used to infer demographic history of a small subset of samples sequenced at high coverage (35-40x). These analyses highlight a historic separation of Haida Gwaii populations with all other populations (**Figure 7**), consistent with past work (McGovern et al., 2010). These results are scaled with a mutation rate of 9.13e^-9^ based on germline mutation rates in the crown-of-thorns sea star (Popovic et al., 2024).

**Figure 7:**
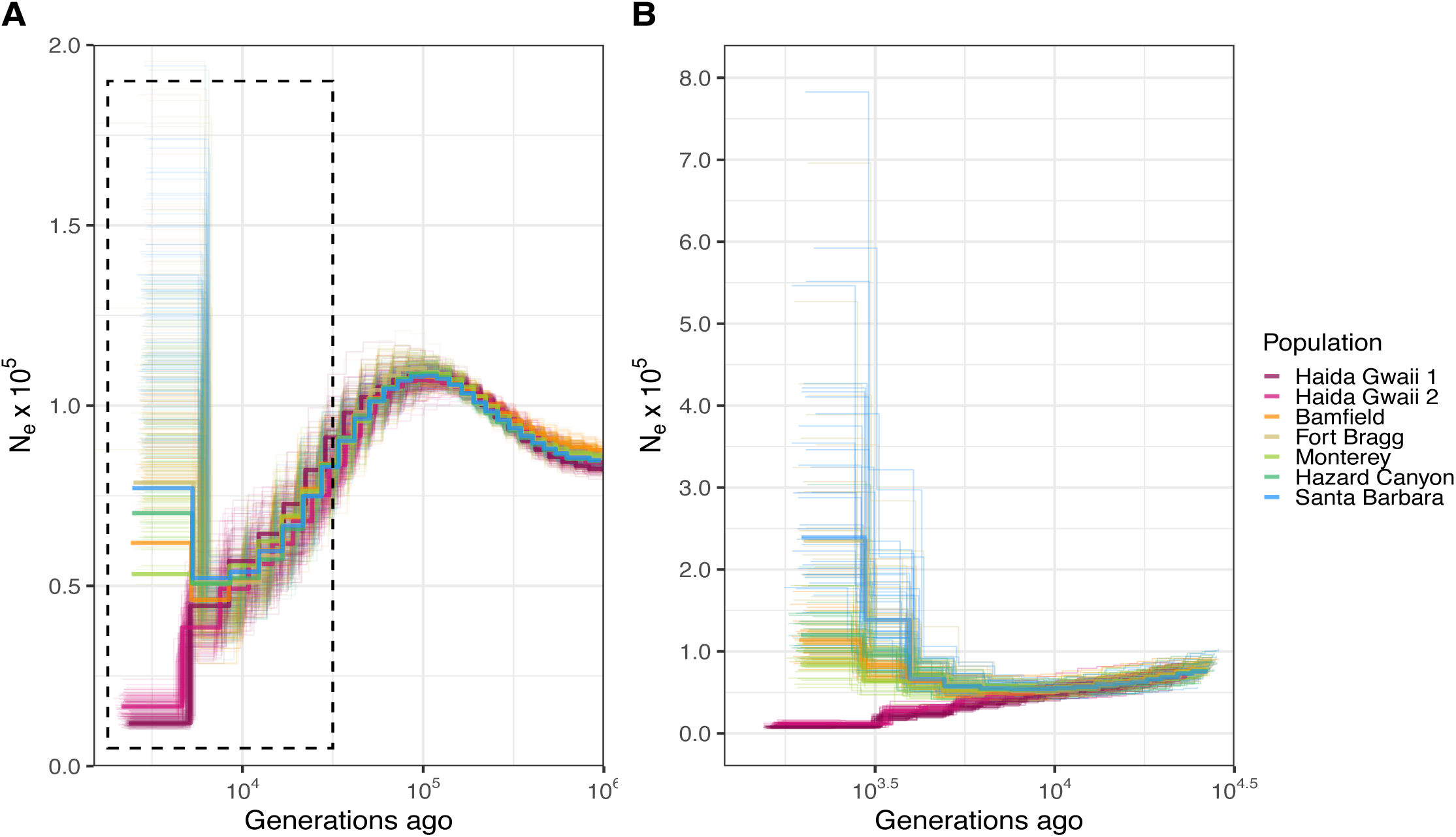
Demographic Analysis. All data in this figure represent analysis of individuals sequenced at high coverage (>30x). Predicted historic effective population sizes (y-axis) are plotted from more recent to older timescales (left to right). PSMC results **(A)** are shown with thin lines showing results from 100 bootstrap replicates. bPSMC results highlight a population bottleneck in Haida Gwaii samples **(B)**. Plot *B* represents a more recent timescale indicated by the dashed box in plot *A*. Light lines in plot *B* indicate results from 40 bootstrap replicates. For both plots, a mutation rate of 9.13e-9 was used.

Central BC populations show the same signature of a population bottleneck. However, this sample and the sample from Ensenada were sequenced at a significantly lower depth (18x). Subsampling our other populations to this lower depth confirms overlap between effective population sizes in all southern samples and similar population bottlenecks in both Central BC and Haida Gwaii populations (**Supp. Figure 2,3**).

### Supplemental Analyses

To assess the likelihood of different values of groups in ngsADMIX, we plotted these likelihoods ins **Supp. Figure 13** (slightly more probable with increasing values of k). To infer the direction of larval dispersal between populations, we used OpenDrift to model larval dispersal in several El Nino and La Nina years (2000, 2010, 2011, 2013, 2016, 2020) over a 10-week period from May 1 - July 10th. While an in-depth analysis of larval dispersal is beyond the scope of this study, modeling highlights that larval dispersal is likely driven by generally southward currents during the summers, strengthening in El Nino years (**Supp. Figure 14**). Treemix analysis similarly infers a general trend of southward dispersal, with the strongest patterns of inferred migration occurring between Haida Gwaii and Central BC. On longer historical timescales, Central BC and Santa Barbara samples are also inferred to have shared migrants. However, this pattern might be driven in part by our Santa Barbara *aquarium* samples (collected around 2008) that both share the highest similarity to our *new* Santa Barbara samples and also retain similarity to northern samples. If these are removed from the analysis, this connection remains to southern samples, but not necessarily to the Santa Barbara population specifically. Again, these results highlight the strong similarity of our southern samples. Nonetheless, we find a fairly strong signature of overall southward movement, consistent with a southward coastal current throughout the sampled range (**Supp. Figure 14,15**).

## Discussion

We describe the population genomics of *P. miniata* and find that this species harbors both highly divergent populations with strong population structure in the northern part of its range and a more panmictic set of populations south of an extensive range gap. Our analyses support high gene flow and reduced spatial genetic structuring among bat star populations throughout California, a large effective population size, and low-moderate nucleotide diversity in this species. We corroborate a previously described strong genetic disjunction across Queen Charlotte Sound (Keever et al., 2009; McGovern et al., 2010; Sunday et al., 2014) and, additionally, identify a much weaker genetic discontinuity spanning the extensive range gap in Washington and Oregon. South of this range disjunction, populations exhibit characteristics of a more panmictic unit. In contrast, northern populations display more pronounced signatures of genetic structuring and isolation. These signatures of structuring and potential local adaptation are likely facilitated by the extensive range gap, as strong selective sweeps in *P. miniata* populations are particularly evident in populations north of the Washington/Oregon range gap.

### Demographic History Shapes Genomic Differentiation in Northern Populations

As previously described by McGovern et al., we find evidence of a genetic split among northern populations, likely aligning with a transition into the last glacial period (Clague et al., 2005; Clague & James, 2002; McGovern et al., 2010) and suggesting persistence of multiple refugia during the last glacial period. PSMC analysis using high coverage genomes in this study suggests considerable population bottlenecks in these northern Haida Gwaii and Central Canadian populations during glaciation, suggesting that these populations experienced considerable population declines during this time. While on long historical timescales, *P. miniata* populations north of Vancouver Island likely experienced population bottlenecks, populations in the southern-most parts of this species’ range likely experienced population expansions, manifesting in diverging population trajectories of these northern and southern populations. On more recent timescales, Treemix analysis and oceanographic modelling (**Supp. Figure 14,15**) suggest that southerly currents during summer months may occasionally bring northern migrants to the southern genetic pool, whereas genetic flow south to north appears less likely. This could be due to the timing of spawning: while *P. miniata* are capable of spawning in the laboratory year-round, peak spawning activity is thought to occur in the field in summer months (Rumrill, 1989; M. F. Strathmann, 1987), coinciding with predominantly southward surface currents (Holt & Mantua, 2009; Mauzole et al., 2020).

### Evidence for Strong Selective Sweeps Across a Range Gap

While glacial cycles likely contributed to population splits in the northernmost populations in this study, we did not find evidence of a demographic split between populations across the range gap. However, we detected strong localized genetic differentiation between these populations. Specifically, we observed high F_ST_ in several genomic loci, many of which were associated with evidence of selective sweeps, particularly in northern populations. Past work has shown that divergence at egg receptor and sperm binding loci may drive isolation of populations across Queen Charlotte Sound (Hart et al., 2014), where the strong genetic disjunction exists. Furthermore, reduced migration across the range gap might limit gene flow from the south to Vancouver Island populations, possibly increasing the probability of changes in allele frequencies in response to environmental pressures. In southern populations there is less evidence of genetic differentiation, aligning with the hypothesis that, in these more interconnected populations, adaptive selection may be swamped by gene flow.

Strong genetic differentiation was detected in genes encoding ion channel proteins, heat shock protein chaperones, and proteins involved in lipid metabolism and glycosylation, as well as a gene associated with mucus production, which shows a strong latitudinal gradient across the range gap (SC 69 in **Figure 5**). In some cases, signatures of selective sweeps overlapped with intergenic loci or proteins involved in transcriptional regulation, highlighting that regulators of gene expression that remain poorly described in many non-model species (Bonasio, 2015) may be important targets of selection. **Supp. Table 3** further describes some of the predicted impacts of SNP differences among populations.

Due to the broad genetic regions which show high levels of differentiation among populations and our lack of knowledge on the function of many proteins in this species, we can only speculate on the function of these genes. One hypothesis for the strong differentiation in genes involved in ion transport, particularly within scaffold 72, might be related to surpassing strong salinity gradients that are a defining feature of water masses in Washington and Oregon, where rivers contribute to reduced salinity. The Columbia River plume, for example, has been shown to block coastal transport of planktonic larvae as this plume contributes to a distinct low salinity water mass (Banas et al., 2009; Brasseale, 2020; Peterson & Peterson, 2009).

As glaciers receded after the last glacial maximum, meltwater could have also contributed to strong riverine outflows contributing to low-salinity water masses along the northern Canadian coastline (Wickert, 2016), possibly creating a strong selective pressure for low-salinity adaptation. The absence of *P. miniata* in Juan de Fuca strait, characterized by low salinities but only modestly different sea surface temperatures, also provides support for this hypothesis. This explanation for our strong selective sweeps could also explain why our Winter Harbor populations have a weaker selective sweep in this region and, overall, share more similarity to southern populations compared to Bamfield or Ucluelet samples, which are geographically closer but experience lower coastal salinities, compared to Winter Harbor (Fatland et al., 2016). However, the absence of this ion-channel associated selective sweep along the Central Canadian coastline is surprising and inconsistent with this hypothesis, since this region is also associated with high riverine outputs. Furthermore, our windowed PCA analysis suggested that the differentiation at this locus was weaker at Fort Bragg populations collected in the past (less than 10 years ago), which could suggest a much more recent divergence at this locus.

It is possible that multiple glacial refuge populations persisted independently, and these glacial refuge populations had historically independent evolutionary trajectories. The Central Canadian population, collected near Bella Bella, is characterized by a high inbreeding coefficient and a unique set of selective sweep loci that is quite diverged from the Haida Gwaii population, despite its relative geographical proximity, suggesting this as a possibility. Considering the high inbreeding coefficients in this Central BC population, it is also possible that this population consists of hybrids between diverged populations north and south of Queen Charlotte sound, supporting past work that has documented lower breeding efficacy of populations across this genetic divergence (Hart et al., 2014; Patiño et al., 2016). Admixture analysis also suggests that this relatively inbred population more closely resembles southern populations, compared to Haida Gwaii, suggesting an evolutionary history that could be distinct from this populations’ northern neighbors.

Notably, the putative selective sweep in scaffold 72 (along with other hypothesized sweep loci) is centered narrowly in the genome, reminiscent of adaptive polymorphic inversions in other marine species. Such divergences are often seen across polymorphic inversions, which can reduce recombination and allow selection on multiple adaptive alleles in ‘super genes’ (Barney et al., 2017; Matschiner et al., 2022; Stenløkk et al., 2022). The geographic patterns shown here, along with patterns in selective sweeps, suggest that these polymorphisms diverged north of the range gap. While we did not find strong evidence of an inversion using a windowed PCA analysis (**Supp.** Figure 5), it is possible that long-read sequencing would be required to resolve this.

In addition to the strong differentiation described within scaffold 72 across the range gap, differentiation of a putative mucin gene with latitude might suggest differing selective pressures associated with mucous production in different thermal regimes, north and south of the range gap. Mucous layers in marine species are composed of glycosylated mucins that can serve to both inhibit and promote binding of bacterial pathogens, depending on their structure (Padra et al., 2019; Van Klinken et al., 1995), which can vary considerably by species (Pajic et al., 2022). Mucins tend to be more effective at high temperatures, where their expression is upregulated in fish (Sanahuja et al., 2019; W. Wang et al., 2022). In echinoderms, these genes are also thought to play an important role in protecting against pathogens (Bavington et al., 2004) and positive selection of mucin in *P. miniata* could be suggestive of different selective pressures in microbial defense. Future functional work to investigate some of the genomic regions in which we find evidence of selection, could shed light on the hypotheses generated in this study.

### No Evidence for Localized Selection before and after Wasting Disease

Northern *P. miniata* samples, including Haida Gwaii, Central Canadian, Winter Harbor, and *historical* Bamfield population were collected and sequenced at the mitochondrial and microsatellite level in the early 2000s by authors from a previous study (Keever et al., 2009; Sunday et al., 2014) and resequenced using lcWGS in this study to identify whether deeper sequencing could identify novel genomic regions under selection and identify possible selection happening after sea star wasting disease, which decimated many sea star species along the Pacific coastline, but left *P. miniata* comparatively unaffected (Dawson et al., 2023; Hewson et al., 2018; Menge et al., 2016; Miner et al., 2018; Montecino-Latorre et al., 2016).

While we observed some considerable variation in the genomics of *P. miniata* collected in different years at localities where more than four samples from before and after this disease were available (Bamfield, Santa Barbara), we do not find evidence for selective sweeps or strong genomic differentiation at specific loci that would be likely to confer disease resilience after this event. However, we hope that this study can provide insight into genomic mechanisms that may have allowed this species to survive this disease, while other species suffered considerable population declines. Recent work on the genomics of *Pisaster ochraceus*, for example, has shown mixed impacts on the genomics of this species, which suffered mass mortalities after wasting disease, highlighting possible shifts in allele frequencies, loss of rare alleles, and increased differentiation among populations following the event (A. R. Burton et al., 2022; Schiebelhut et al., 2018, 2022). Our results indicate a similar, though modest, trend of decreasing rare alleles (increased Tajima’s D) in *new* samples from Santa Barbara, but no difference in Bamfield between *new* and *historical* samples. In the case of *Pisaster ochraceus*, Schiebelhut et al. proposed that the observed reduction in rare alleles may have resulted from genetic drift in reduced population sizes, potentially diminishing the species’ adaptive capacity in future generations. While, in *P. miniata* mortalities were not as common, it is possible that some populations still lost rare variants. Comparative genomic work on multiple Pacific coast sea star species differentially impacted by the wasting disease, could shed valuable light on the genomic signatures of sea star wasting disease on different echinoderm species.

### Sex Differences

One surprising finding of this work is the strong signature of male/female differences in our overall visualizations of population structure (i.e. **Figure 2** and **Supp. Figure 1**). This divergence is most clear when looking at populations south of the range gap. Here, this divergence between males and females is driven primarily by two neighboring genomic scaffolds (**Supp. Figures 4,5**). Some of our analyses suggested that these regions may correspond to a sex chromosome, as male and female versions of this extended region do not seem to recombine (**Supp. Figure 5)**. However, SNP heterozygosities are high in both males and females and sample coverages in males are not as high as would be expected following this hypothesis (only slightly higher in males). These results could be obtained if an additional female chromosome (i.e. in a ZW sex-determination system) shares homology to the male sex chromosome and aligns quite well to male-version scaffolds. While *P. miniata* is known to have distinct sexes, other asteroid species are known to change sex or be hermaphroditic (Ebert, 2021; R. R. Strathmann et al., 1984) suggesting some flexibility in sex determination system within the class. Previous work in molluscs has highlighted that longstanding conservation of similarity between female ZW and male Z chromosomes can occur in marine invertebrate species, with enhancer translocation between chromosomes providing a mechanism for modifying sex determination between closely related species (Han et al., 2022). A similar system in asteroids could account for many of the observations in this work and differences in sex-determination within asteroids. However, mechanistic studies would be needed to further investigate this. Nonetheless, we provide strong evidence of a sex-determining region in two scaffolds that differ between males and females in *P. miniata*. Data and understanding of sex-based differences in population and community structure and dynamics, and responses to climate change, is limited in marine and terrestrial systems, but review and syntheses of available studies indicate significant variation, highlighting this is an important area for future research (Gissi et al. 2023a,b).

### *P. miniata* Population Genomics in Context

Our finding that strong differentiation of specific genetic loci and selective sweeps within populations in *P. miniata* highlights that important differentiation can occur within marine invertebrate populations even in the absence of strong genetic structuring and presence of a long-lived planktonic larval period (Barney et al., 2017; Lee et al., 2024; Oliver et al., 2010; Reeb & Avise, 1990; Sanford & Kelly, 2011; H. Wang et al., 2024). Notably, we see some genomic differences within nearby populations between sampling years, which could be suggestive of year-to-year differences in the genetic pools that contribute to local populations, as has been hypothesized for many other marine invertebrate species with planktonic larvae (Carr et al., 2003; Rumrill, 1989). However, as we only have two temporal time points for a limited number of populations, this remains speculative. Despite this limitation, this study offers a rare opportunity to analyze temporal variation in sampling. Our results highlight the need for studies that further examine how fine-scale genetic patterns can vary between years and on decadal time-scales, within a species. Such studies could greatly improve our understanding of how larvae disperse and settle in the ocean and how year-to-year variability in environmental conditions and oceanography impact this, tying together theoretical work on dispersal (Lett et al., 2010; Swearer et al., 2019) with observations of genetic patchiness and high variability in year-to-year recruitment for many marine invertebrate species (Botsford, 2001; Broitman et al., 2008; Fogarty et al., 1991; Selkoe et al., 2010).

In other broadly dispersing marine invertebrate species, some studies have found very little genetic structure while others observe high genomic differentiation in different locations along the Pacific coastline (R. S. Burton, 1997; Griffiths et al., 2025; Kelly & Palumbi, 2010, p. 50; Palumbi, 1994, 1995; Sanford & Kelly, 2011). Similarly, while some studies have observed that life history strategies and larval durations can sometimes be important predictors of genetic structure (Bingham, 1992; Ross et al., 2009; Schinske et al., 2010; Sherman et al., 2008; Torrado et al., 2021), many studies and reviews do not observe clear trends in which taxa or life history strategies are more likely to be associated with strong genetic structure or spatial disjunctions (Arranz et al., 2021; Costantini et al., 2018; Faurby & Barber, 2012; Ludt & Myers, 2021; Riginos et al., 2011; Selkoe & Toonen, 2011), highlighting that complex factors can shape marine population dynamics. In *P. miniata*, it seems likely that historic isolation of populations during glacial periods allowed for isolation and selection (McGovern et al., 2010) while oceanographic features, like the North Pacific current that splits along the Canadian coastline (Keever et al., 2009; Tanhua et al., 2013; Wells et al., 2008) and riverine outflows that impact water masses near Washington and Oregon (Banas et al., 2009; Brasseale, 2020; Peterson & Peterson, 2009), along with reproductive barriers (Hart et al., 2014; Patiño et al., 2016) could contribute to maintaining strong population divergence in northern populations.

This work highlights that, for marine invertebrate species, which often have large effective population sizes and high inter-population gene flow (Carr et al., 2003), genomic analyses may still detect structuring between subpopulations where microsatellite or mitochondrial data are unable to resolve these. Furthermore, this work demonstrates that relationships between genetic structure and larval dispersal, where high dispersal potential has been associated with decreased genetic structure (Schiebelhut & Dawson, 2018), can be highly context-dependent (Weersing & Toonen, 2009).

The reduction in sequencing costs in recent years makes large-scale genomic analysis in coastal marine species much more feasible than they have been previously (Lou et al., 2021). In the context of climate change and large-scale anthropogenic habitat destruction, effectively planning and prioritizing areas for marine and coastal protection is becoming increasingly urgent, and genomic characterizations of marine species are increasingly being considered when deciding on measures for effective conservation (Jeffery et al., 2022). This study sheds light on the complex dynamics of population genetics at sea, using *P. miniata* as a model for understanding marine invertebrates with sedentary adults and long-lived larval stages. In the context of marine conservation, these findings support networks of small Marine Protected Areas (MPAs), informed by evolutionary history and strategically distributed to safeguard both source populations, which replenish neighboring areas, and peripheral populations that can harbor unique genetic diversity (Carr et al., 2003; Ciotir et al., 2013; Jeffery et al., 2022; Lesica & Allendorf, 1995).

## Conclusion

In this study, we find that, despite the high genetic similarity of *P. miniata* along the Pacific coastline south of a known genetic disjunction, strong signatures of local selective sweeps are evident. These findings highlight the occurrence of strong selection and local adaptation in marine species, even in the presence of considerable gene flow. Our results underscore the complex interplay between gene flow, evolutionary history, oceanography, and selective pressures in shaping the genetic landscape of marine organisms and present genomic approaches for detecting selection and local adaptation.

## Ethics and Integrity Statements

### Data availability statement

Scripts for all filtering and analysis steps are available [here](https://github.com/vpagowski/Pminiata_popgen_paper.git). All sequence data and sample metadata have been submitted to the SRA under project ID PRJNA1269039.

## Acknowledgements

We would like to thank multiple aquaria that donated samples to this project including the Noyo Center for Marine Science, Santa Barbara Sea Center, Heal the Bay Aquarium in Los Angeles, and Birch Aquarium in San Diego. Thank you as well to Dr. Michael Hart for donating historical samples to this study. Likewise, we would like to thank Bamfield Marine Sciences Center staff for coordinating collections in Bamfield. Thank you to friends and colleagues, including Melissa Palmisciano, who helped collect samples. We would also like to thank R. Beas-Luna for providing the samples from Baja California and *Texas A&M AgriLife Research: Genomics and Bioinformatics Service* for library preparation and sequencing of samples from this study.

Finally, we would like to acknowledge support from NSF (2108566 to FM), as well as the Irene Brown and Friends of Hopkins graduate research grants to VP.

## Benefit Sharing

All collaborators who contributed to project conception, analysis, or guidance are included as co-authors and the results of research have been shared with the groups and organizations that provided samples. More broadly, our group is committed to international scientific partnerships, as well as institutional capacity building and we welcome the use and sharing of any of the results or data from this project.

## Author Contributions

VP, SP, and FM conceived and designed the study. VP analyzed sequencing data and wrote the original manuscript. EH and BC provided valuable input on analyses, pipelines, and provided feedback on the original manuscript, along with SP and FM.

