## Supplemental Figures and Tables for "Population Genomics of *P. miniata* along the Pacific Coastline Reveal Subtly Diverging Genomics Along an Extensive Range Gap"

### Supplemental Tables

**Table S1:** Sample information for individuals used in this study

| Location | Approx Lat | Approx Long | Collection | Collection Time | Number | Sequencing Round | Notes |
|---|---|---|---|---|---|---|---|
| Haida Gwaii | 52.96 | -132.43 | Historic | 2005-Jun | 12 | 1 | From Hart Lab |
| Central BC | 51.88 | -127.98 | Historic | 2010-Sep | 10 | 1 | From Hart Lab |
| Winter Harbor | 50.46 | -128.07 | Historic | 2006-Aug | 12 | 1 | From Hart Lab |
| Ucluelet | 49 | -126 | New | 2023-Sep | 11 | 1 | |
| Bamfield (Scott's Bay) | 48.89 | -125.11 | New | 2023-Sep | 5 | 1 | Grouped in analysis |
| Bamfield (Aguilar Point) | 48.89 | -125.11 | New | 2023-Sep | 17 | 1 | |
| Bamfield (Aguilar Point) | 48.89 | -125.11 | Historic | 2006-Jan | 10 | 2 | Grouped in analysis |
| Bamfield (Scott's Bay) | 48.89 | -125.11 | Historic | 2004-Nov | 10 | 2 | |
| Fort Bragg | 39.43 | -123.81 | New | 2023-May | 28 | 1 | |
| Fort Bragg | 39.43 | -123.81 | Aquarium | 2016-2017 | 10 | 1 | From Noyo Center for Marine Science |
| Fort Bragg | 39.43 | -123.81 | Historic | 2005-Jul | 4 | 1 | From Hart Lab |
| Monterey | 36.63 | -121.94 | New | 2023-May | 28 | 1 | From Monterey Abalone Co. |
| Hazard Canyon | 35.29 | -120.88 | New | 2023-Jun | 11 | 1 | |
| Santa Barbara | 34.42 | -119.64 | New | 2023-Sep | 12 | 1 | |
| Santa Barbara | 34.42 | -119.64 | Aquarium | Likely 2008 | 15 | 1 | From Santa Barbara Sea Center |
| LA | 33.74 | -118.41 | Aquarium | Unknown, likely up to 10 years ago | 15 | 1 | From Heal the Bay Aquarium |
| San Diego | 32.86 | -117.256 | Aquarium | 2016-Feb | 8 | 2 | From Birch Aquarium |
| San Diego | 32.86 | -117.256 | Historic | 2005-Jul | 4 | 2 | From Hart Lab |
| Baja | 31.701 | -116.684 | New | 2023-Oct | 16 | 2 | |

**Table S2:** Filtering summary and number of variants retained for each analysis

| Round 1 | | |
|---|---|---|
| **Total changes** | 49,308,015 | **Percentage** |
| **SNPs** | 41,093,569 | 83.34% |
| **Indels** | 11,806,992 | 23.95% |
| **Multiallelic SNPs** | 2,898,578 | 5.88% |
| **Included** | 9316635 | **18.89%** |
| **Round 2** | | |
| **Total changes** | 75307258 | **Percentage** |
| **SNPs** | 61518113 | 81.69% |
| **Indels** | 17690779 | 23.49% |
| **Multiallelic SNPs** | 3341474 | 4.44% |
| **Included** | 9742686 | **12.94%** |
| **Combined SNPs (both rounds) after filtering** | 8338770 | **16.91%** |

| **Imputed data** | | **Percentage** |
|---|---|---|
| **Total sites** | 581152781 | |
| **Total SNPs after filtering** | 30008952 | **5.16%** |
| **Total SNPs after filtering** Filtering for SNPs present in at least 10 individuals (Figure 2D) | 3804220 | **6.55%** |

**Supplemental Table 3:** For 4 loci outlined in main figure 4, corresponding outputs from snpEff documenting the total, length, number of SNPs, avg. number of bases between variants (variants rate), and the number of predicted impacts on gene function (high, moderate, low). Modifier impacts refer to SNPs outside genes.

**SC55:**

| Chromosome | Length | Variants | Variants rate |
|---|---|---|---|
| **NW_024037055.1** | 15,691,589 | 879 | 17,851 |
| **Total** | 15,691,589 | 879 | 17,851 |

| Type | Count | Percent |
|---|---|---|
| **HIGH** | 1 | 0.031% |
| **LOW** | 499 | 15.373% |
| **MODERATE** | 267 | 8.226% |
| **MODIFIER** | 2,479 | 76.371% |

**High/Moderate Impact:** *Multiple uncharacterized loci (236 variants)*

**SC68 (r1)**

| Chromosome | Length | Variants | Variants rate |
|---|---|---|---|
| **NW_024037068.1** | 28,030,815 | 176 | 159,265 |
| **Total** | 28,030,815 | 176 | 159,265 |

| Type | Count | Percent |
|---|---|---|
| **LOW** | 41 | 4.08% |
| **MODERATE** | 4 | 0.398% |
| **MODIFIER** | 960 | 95.522% |

**High/Moderate Impact:** *Unchar. WD repeat containing protein (1 variant), octopamine receptor 1 (1 variant)*

**SC68 (r2)**

| Chromosome | Length | Variants | Variants rate |
|---|---|---|---|
| **NW_024037068.1** | 28,030,815 | 100 | 280,308 |
| **Total** | 28,030,815 | 100 | 280,308 |

| Type | Count | Percent |
|---|---|---|
| **LOW** | 4 | 1.176% |
| **MODERATE** | 1 | 0.294% |
| **MODIFIER** | 335 | 98.529% |

**High/Moderate Impact:** *Adenosine receptor A2b (1 variant)*

**SC69**

| Chromosome | Length | Variants | Variants rate |
|---|---|---|---|
| **NW_024037069.1** | 46,651,055 | 630 | 74,049 |
| **Total** | 46,651,055 | 630 | 74,049 |

| Type | Count | Percent |
|---|---|---|
| **LOW** | 2,078 | 8.941% |
| **MODERATE** | 1,242 | 5.344% |
| **MODIFIER** | 19,922 | 85.716% |

**High/Moderate Impact:** *Mucin 17/ mucin 19 (73 variants), zinc finger protein 667 (7 variants)*

**SC72**

| Chromosome | Length | Variants | Variants rate |
|---|---|---|---|
| **NW_024037072.1** | 41,642,996 | 208 | 200,206 |
| **Total** | 41,642,996 | 208 | 200,206 |

| Type | Count | Percent |
|---|---|---|
| **LOW** | 8 | 2.395% |
| **MODERATE** | 1 | 0.299% |
| **MODIFIER** | 325 | 97.305% |

**High/Moderate Impact:** *Sodium-calcium exchanger 3 (1 variant)*

### Supplemental Figures

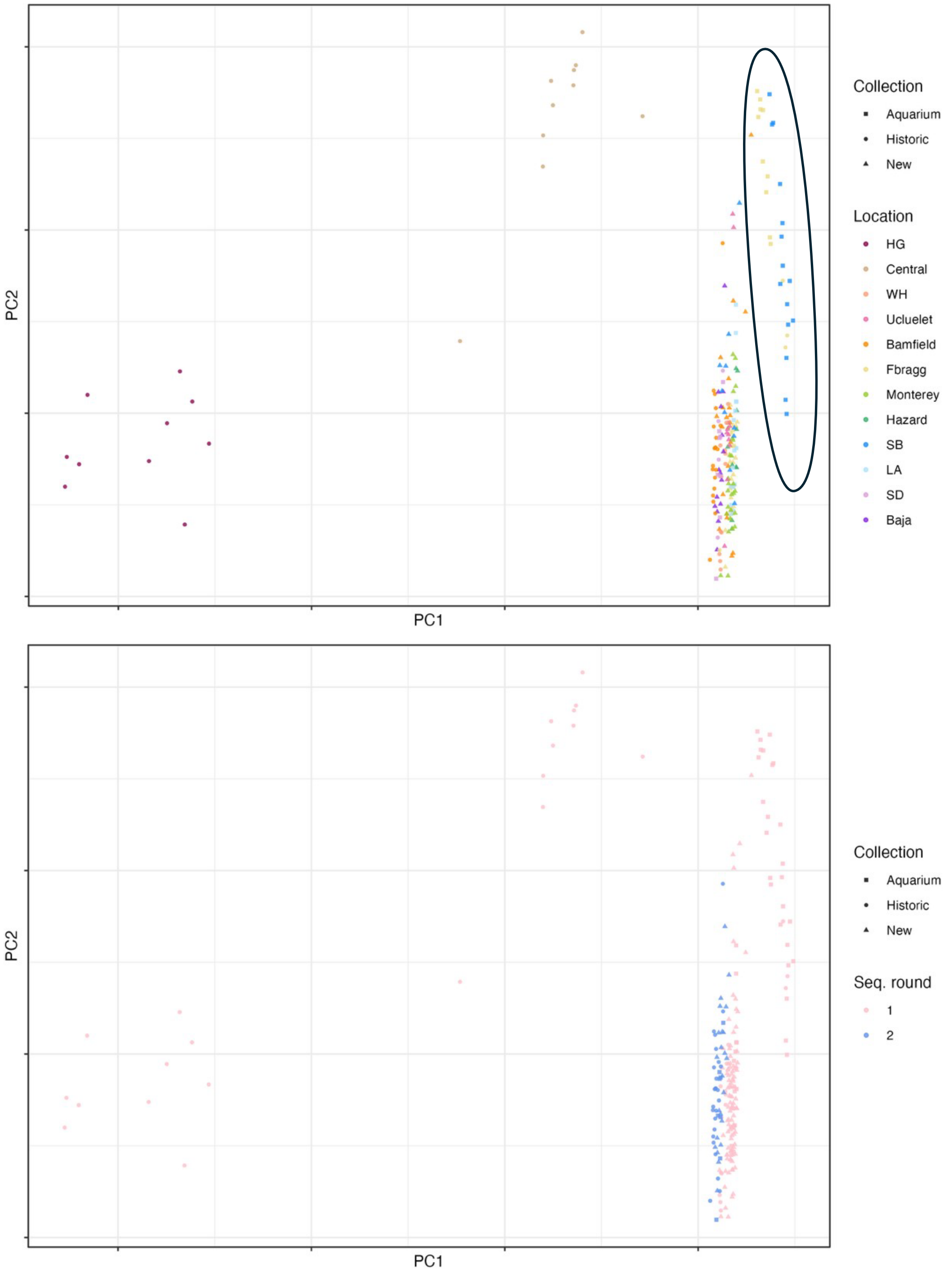


**Figure S1:** PCA from hard call genotypes showing a strong split across Queen Charlotte Sound (two populations furthest to the left on PC1) and clustering of *aquarium* samples from Fort Bragg and Santa Barbara, along with one *new* sample from Bamfield and *historic* samples from Fort Bragg (circled). PCs from this PCA were used to compute covariates in GWAS results from hard call SNPs. Bottom plot shows the sequencing round and batch artefact in the hard call PCA.

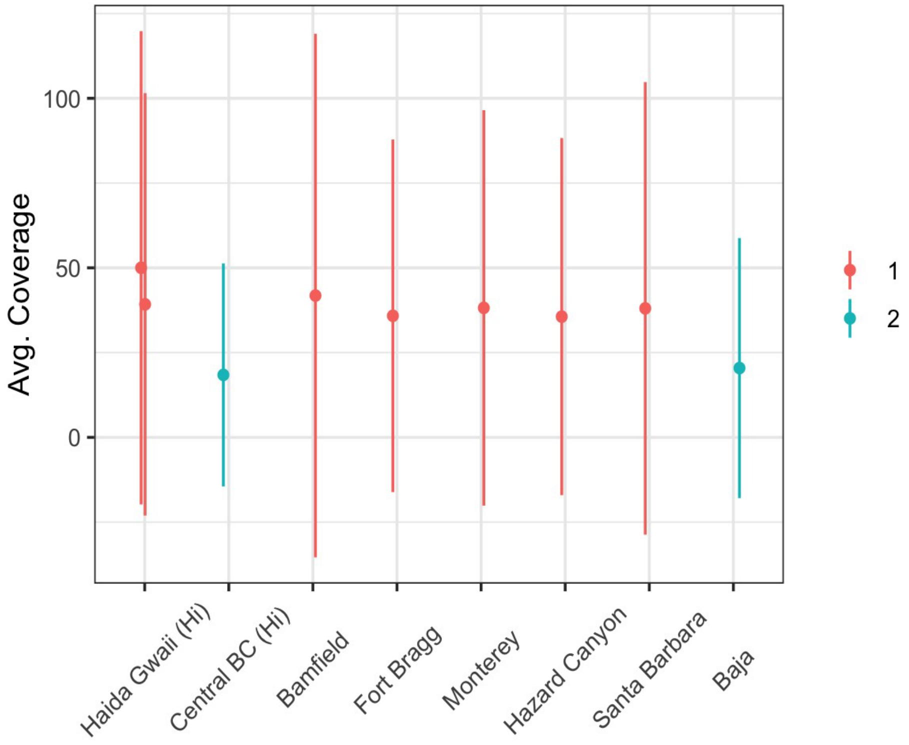


**Figure S2:** Average coverage and standard deviation by location of high coverage samples, with colors indicating which sequencing round samples were sequenced in.

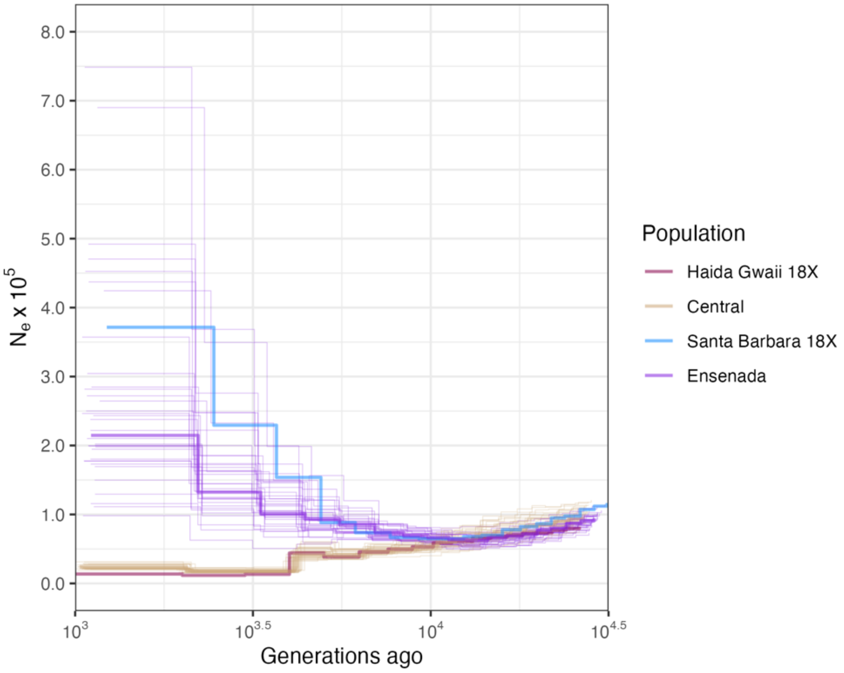


**Figure S3:** Beta PSMC results including Central BC and Baja populations, with our Santa Barbara sample subsampled top 18x suggest a similar expansion of Baja and Santa Barbara populations, using a mutation rate from the crown of thorns starfish (9.13 e-9)

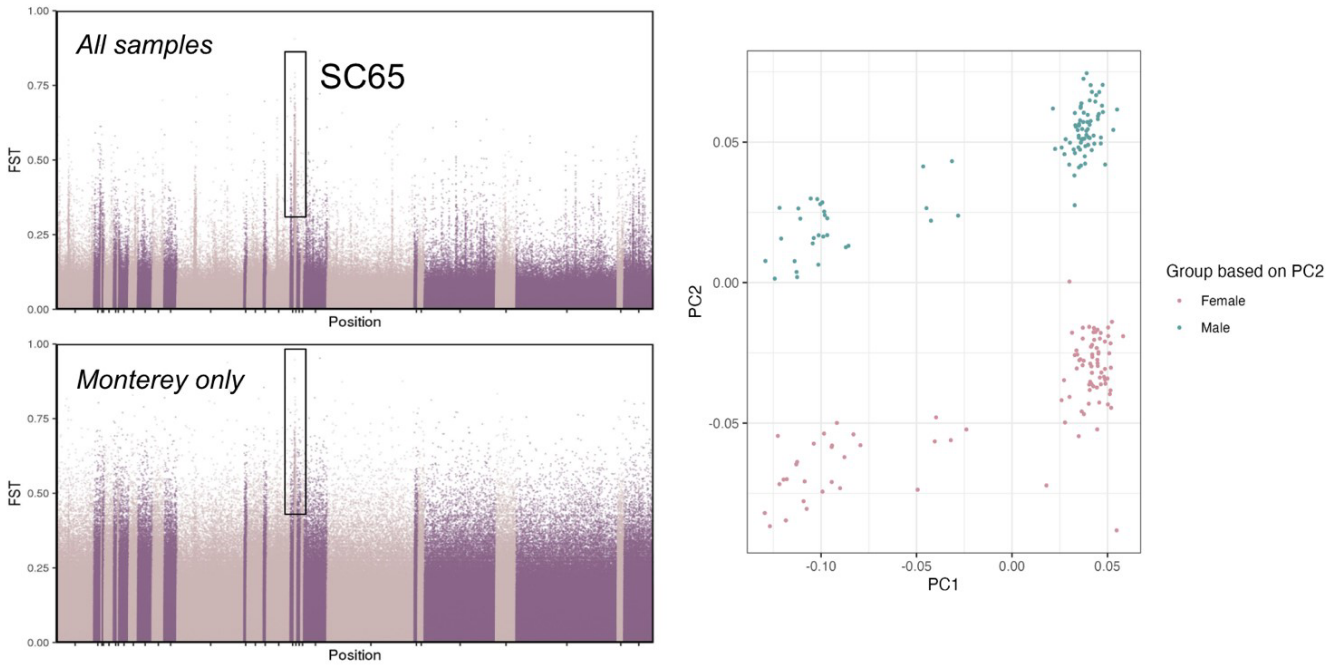


**Figure S4:** Evidence for sex differences between hypothesized male and female samples by categorizing samples based on their position along PC2 from Figure 2 (main figures). When all samples are included or just samples from Monterey (where we had 10 sexed individuals), SC 65 shows higher Fst.

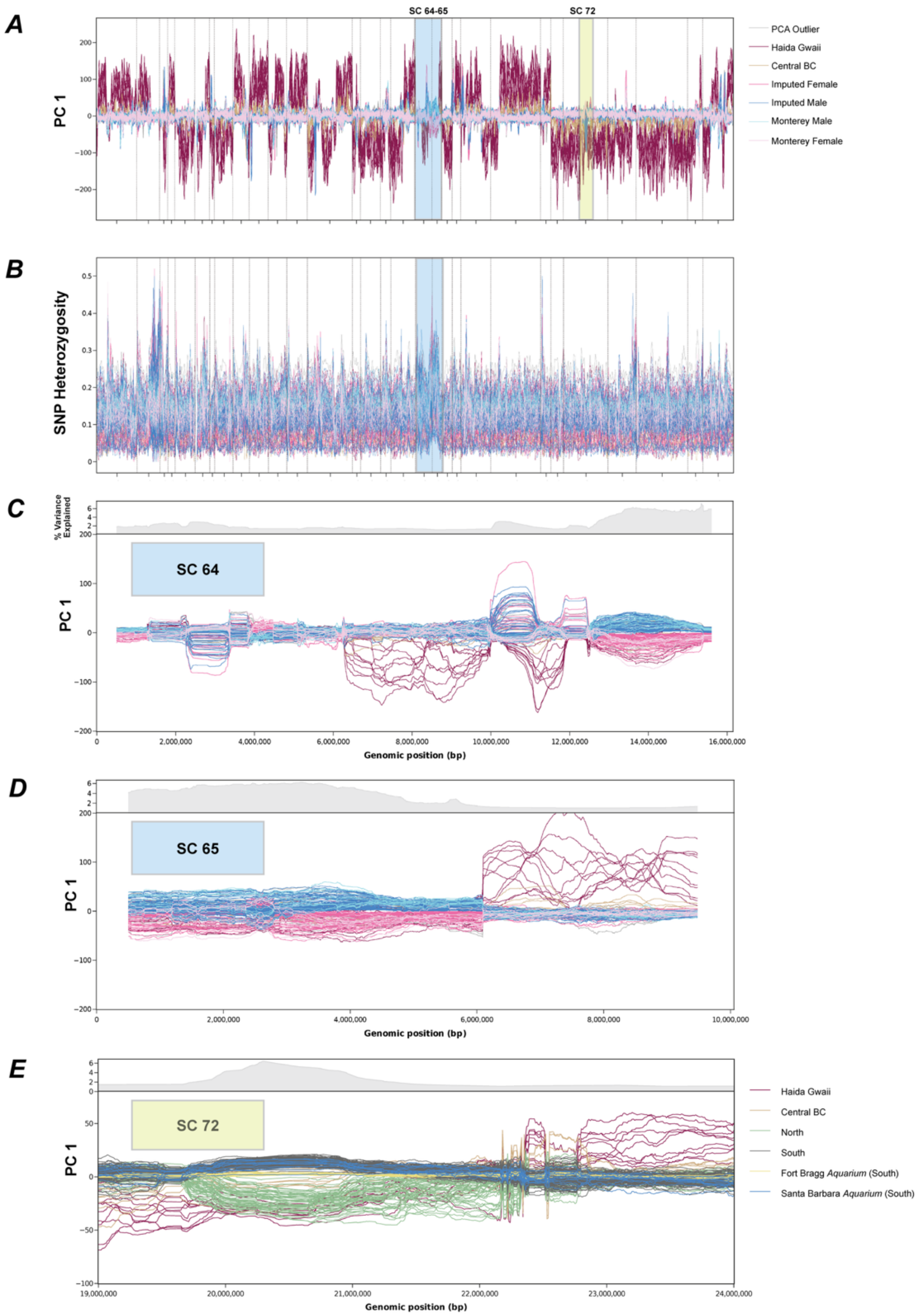


**Figure S5:** Genome-wide windowed PCA results **(A)** and SNP heterozygosity **(B)** with scaffolds 64-65 highlighted in blue and scaffold 72 highlighted in yellow. These scaffolds represent sex-differentiated regions, and a region differentiated across the range gap, respectively. PCA outlier lines plotted in grey represent unsexed individuals (no assignment for imputed sex based on PCA results from **Figure S4** above). A windowed PCA analysis of individual scaffolds for the two sex-associated regions **(C and D)**, suggests lack of recombination throughout most of these scaffolds, between males and females. A windowed PCA analysis of a subset of scaffold 72 shows differentiation of individuals north and south of the range gap, with some *aquarium* samples from Fort Bragg and Santa Barbara showing intermediate signatures of differentiation at the genomic region **(E)**. Y axes for each plot represent PC1 eigenvalues (bottom) and percentage of variance explained by PC1 along the region of interest. Plots were generated with WinPCA software.

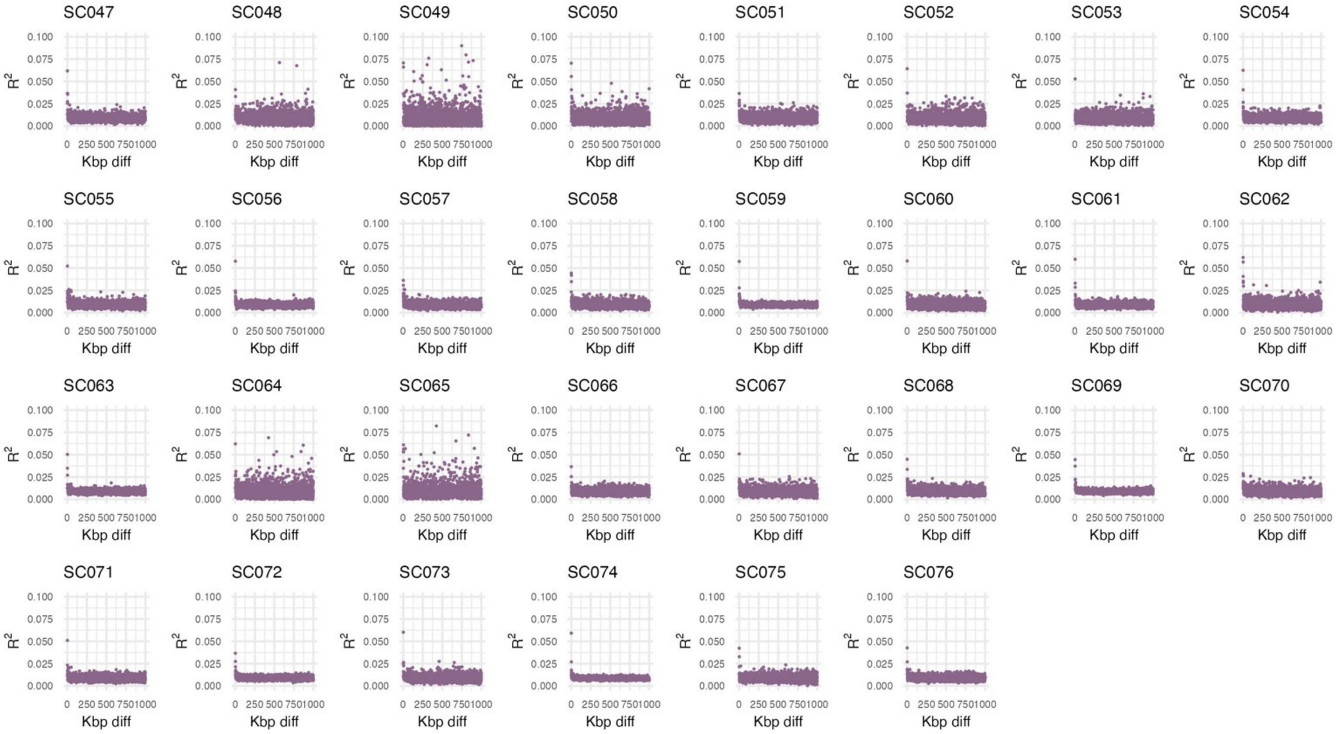


**Figure S6:** Linkage disequilibrium by scaffold, after linkage pruning. R^2 values are higher when linkage between SNPs is higher. The X axis indicates the distance (in kilobases) between SNPs compared. Linked SNPs remain throughout scaffolds 64 and 65 (associated with sex-differences)

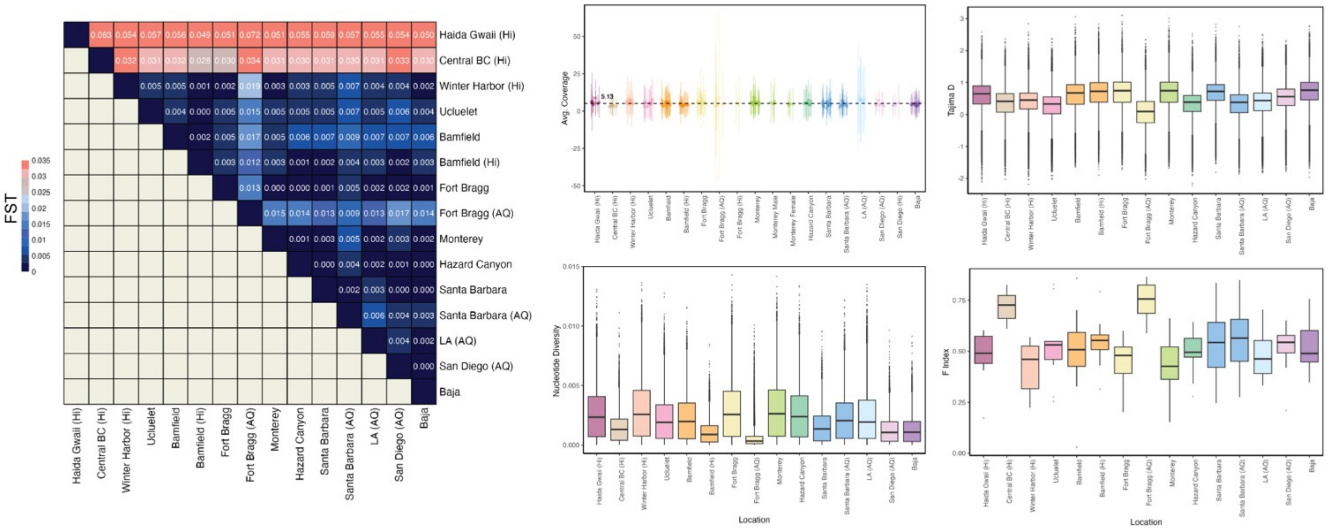


**Figure S7:** This figure is analogous to main figure 3, but showing *aquarium* samples from Fort Bragg

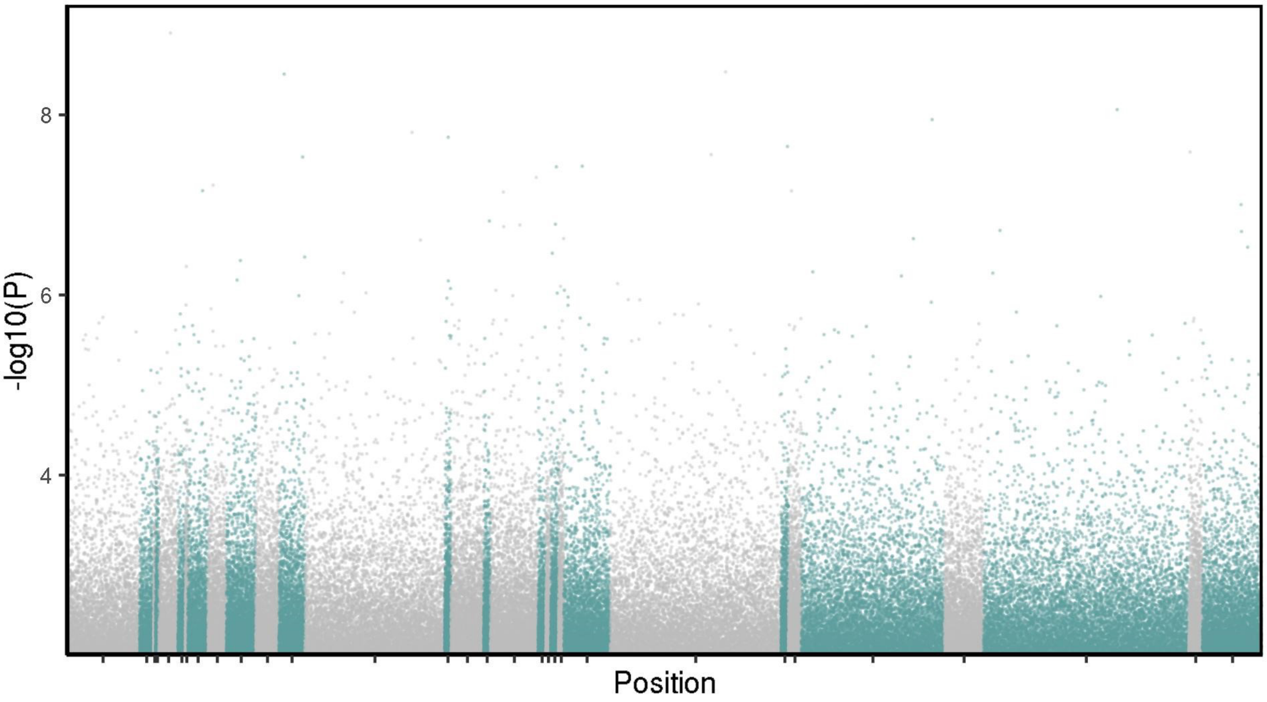


**Figure S8:** GWAS results (using collection latitude as the response variable) excluding all samples north of the range gap shows no evidence of differentiated peaks.

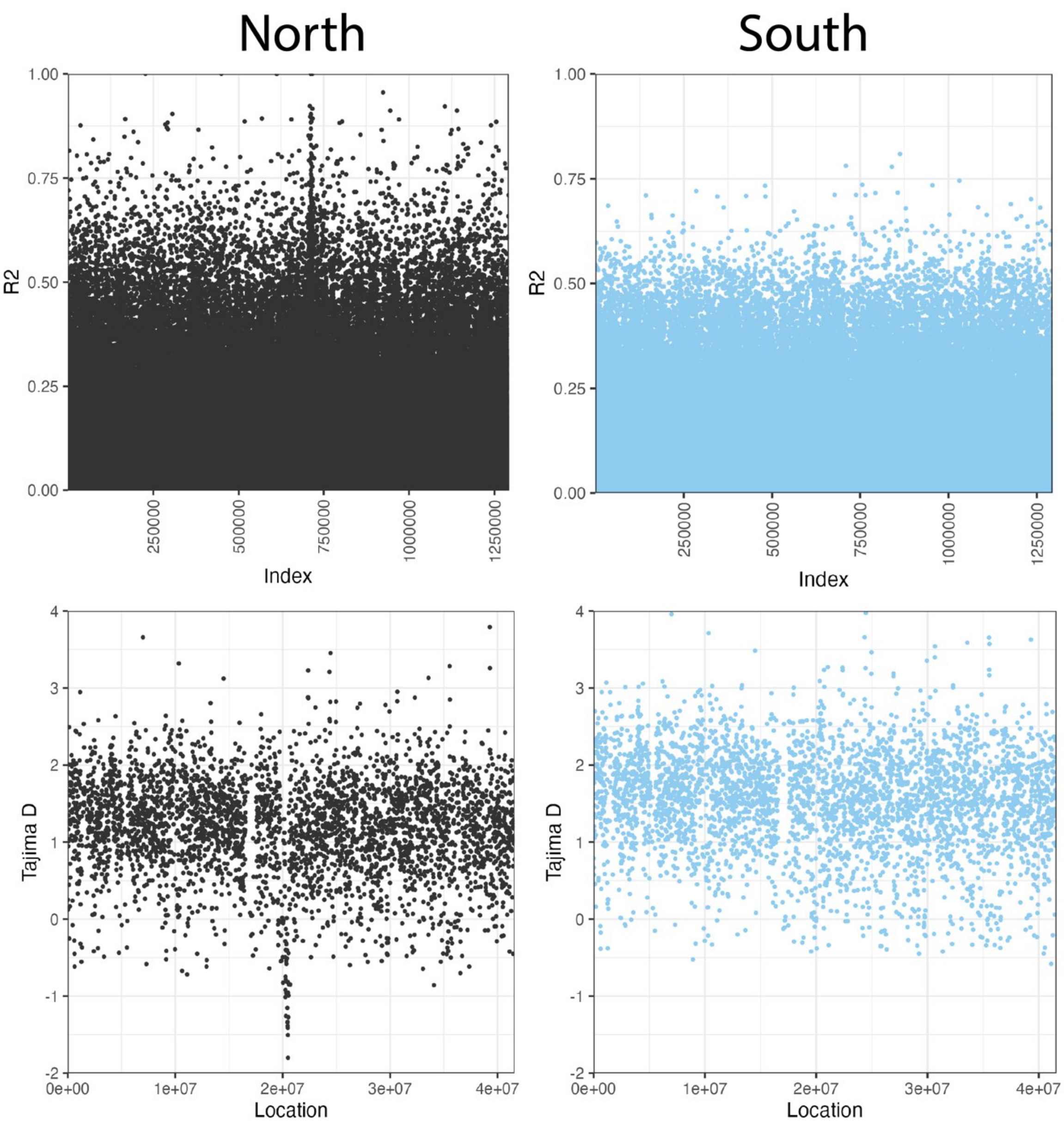


**Figure S9*:*** Pairwise linkage disequilibrium R^2 values (top row) and Tajima's D (bottom row) for SC 72 in the location of the strong selective sweep. Note that the location of high linkage disequilibrium matches the location of the sweep, but the index will be different since these values represent comparisons among SNPs. Northern samples (all samples north of the range gap) are plotted in black and southern samples (all samples south of the range gap) and plotted in blue.

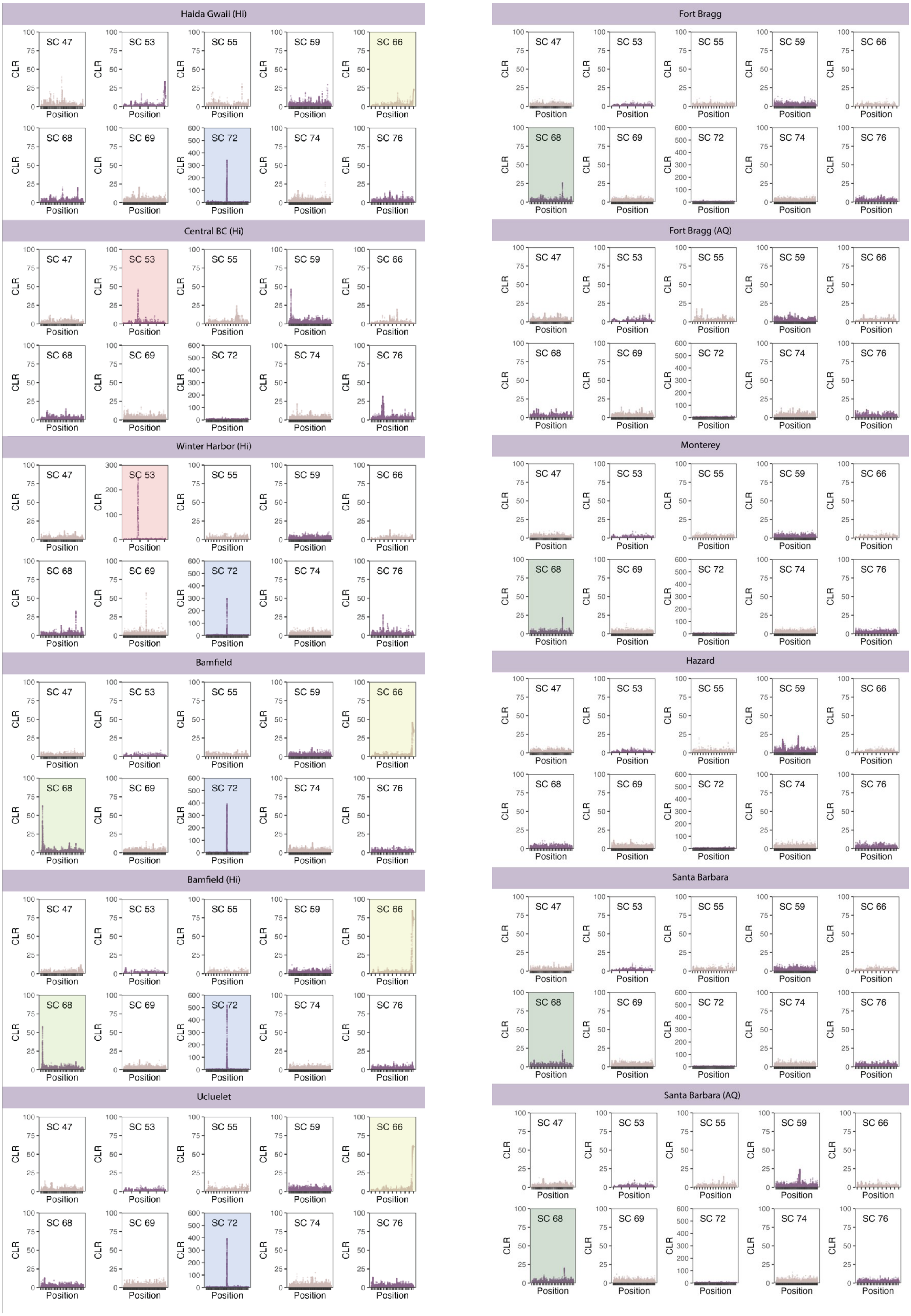

Haida Gwaii (Hi)
Central BC (Hi)
Winter Harbor (Hi)
Bamfield
Bamfield (Hi)
Ucluelet
Fort Bragg
Fort Bragg (AQ)
Monterey
Hazard
Santa Barbara
Santa Barbara (AQ)
SC 47
SC 53
SC 55
SC 59
SC 66
SC 68
SC 69
SC 72
SC 74
SC 76
CLR
Position

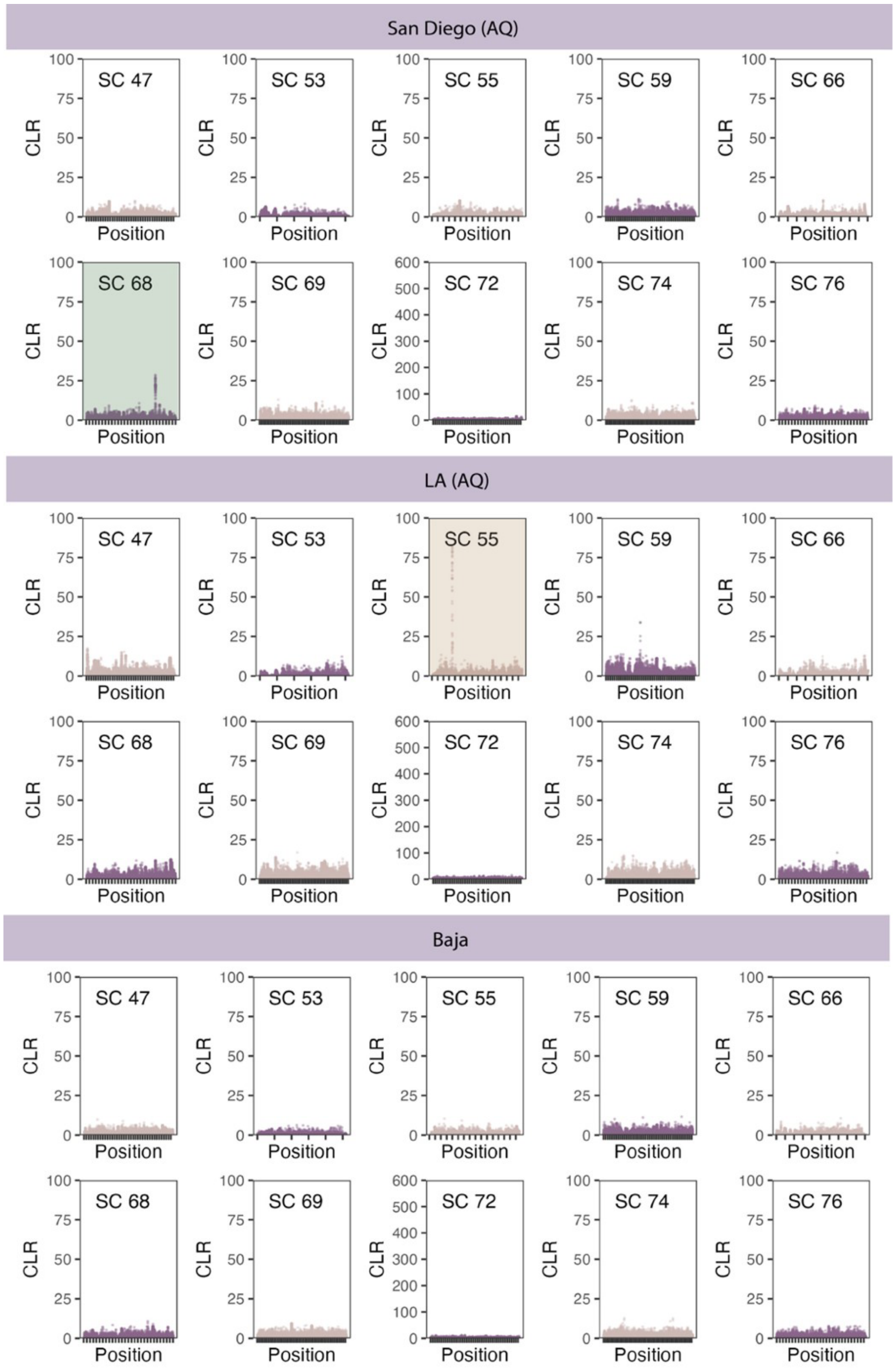


**Figure S10*:*** Selective sweeps in scaffolds of interest. From north to south, starting with the previous page, these figures show cumulative likelihood ratios (y axis, CLR) for selective sweeps calculated with SweepFinder2 for 10 locations of interest. Highlighted plots of the same color indicate the same sweep in different populations. For northern populations, each of these highlighted sweeps has a CLR of at least 75 in one population. In southern populations, each of these highlighted sweeps has a CLR of at least 25 in one population. Tick marks indicate increments of 1 Mbp.

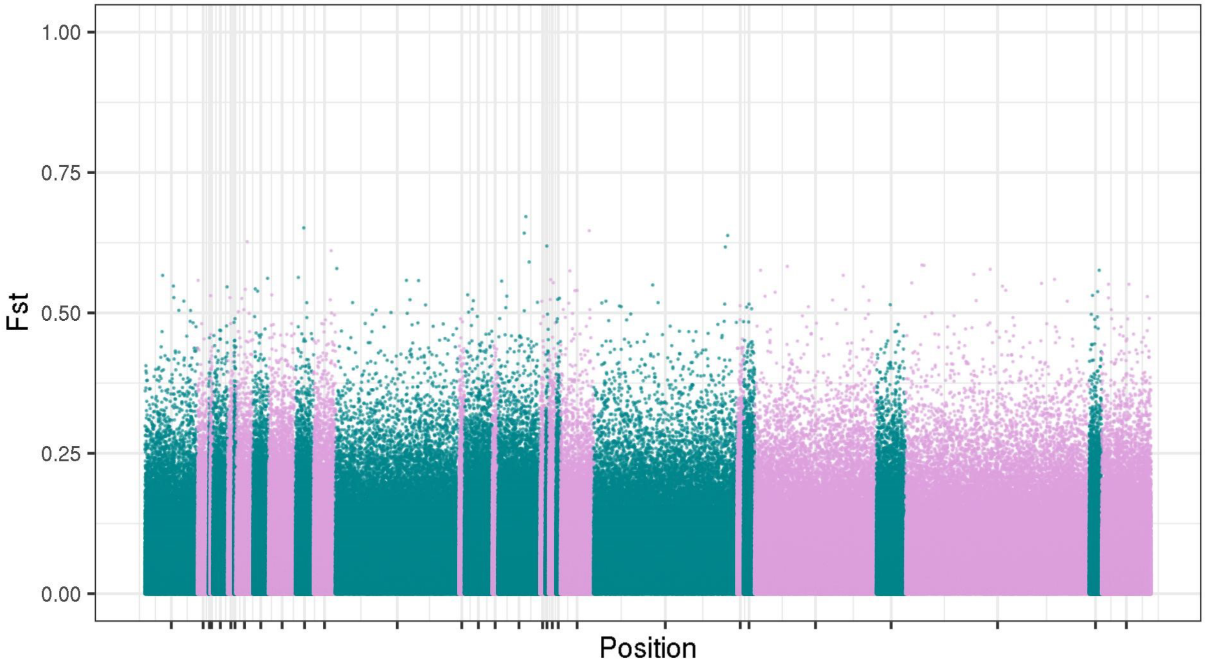


**Figure S11*:*** Pairwise Fst calculated by site for *historic* vs *new* Bamfield samples

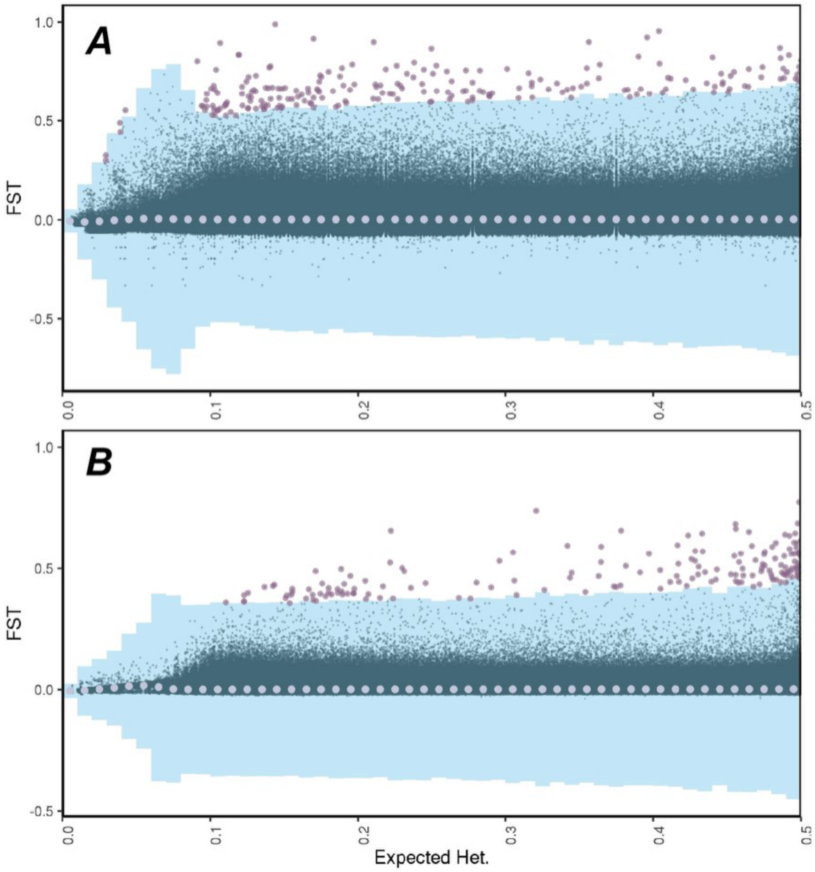


| Go Term | P value |
|---|---|
| Binding | 3.00E-06 |
| Molecular function | 0.0001 |
| Ion binding | 0.0002 |
| Small molecule binding | 0.0004 |
| Adenyl nucleotide binding | 0.008 |

| Go Term | P value |
|---|---|
| calcium:monoatomic cation antiporter activity | 0.002 |
| calcium:sodium antiporter activity | 0.002 |

**Figure S12*:*** Fst vs expected heterozygosity for the entire genome of *P. miniata.* Fst represents comparisons north and south of the range gap, excluding Haida Gwaii and Central BC samples. Blue bars indicate 20 standard deviations from the mean of each interval (increments of 0.01), shown with light purple circles. Outliers are plotted in dark purple after filtering for up to 50% missingness **(A)** and up to 30% missingness **(B)**. Tables to the right represent GO terms identified from these outliers with *p* values under 0.01. GO terms were identified with https://biit.cs.ut.ee/gprofiler/gost, searching for *all known genes*.

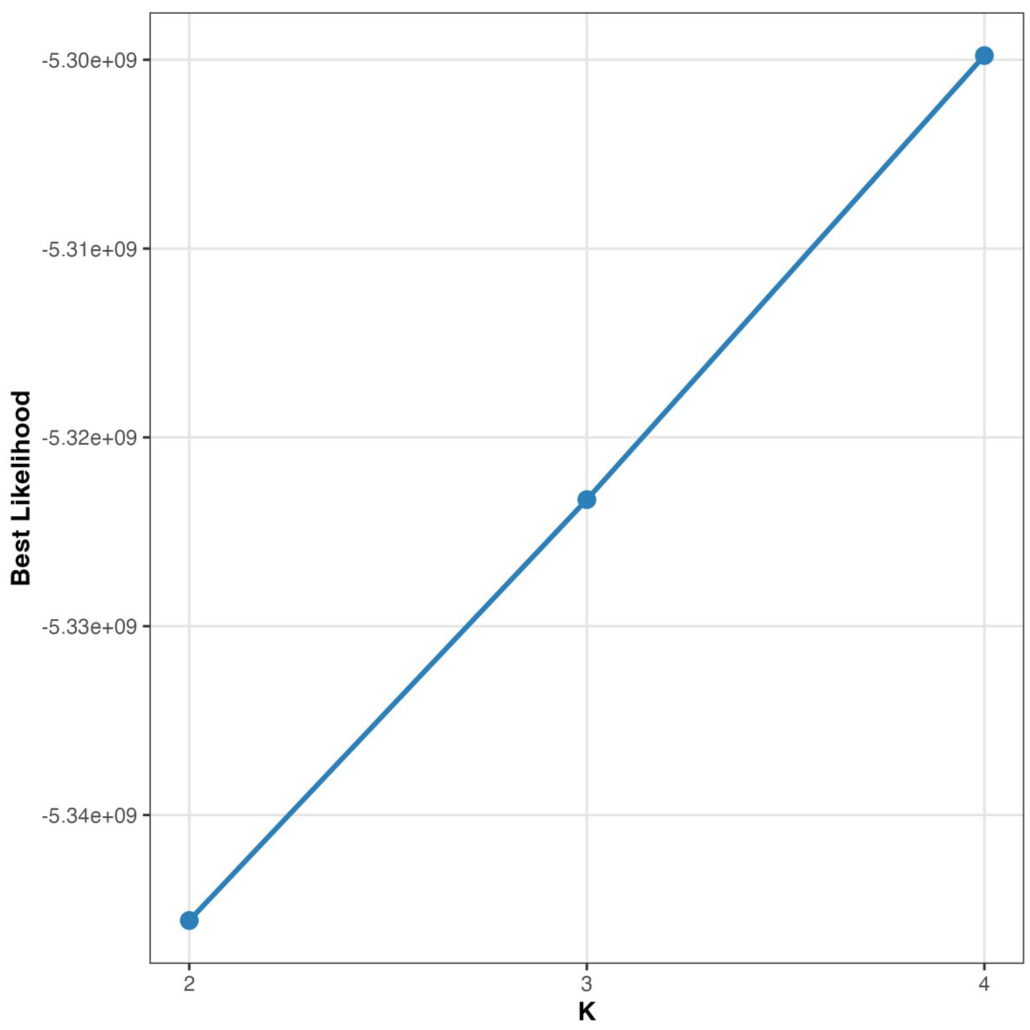


**Figure S13:** Likelihood values from ngsAdmix analysis using k=2-4 groups

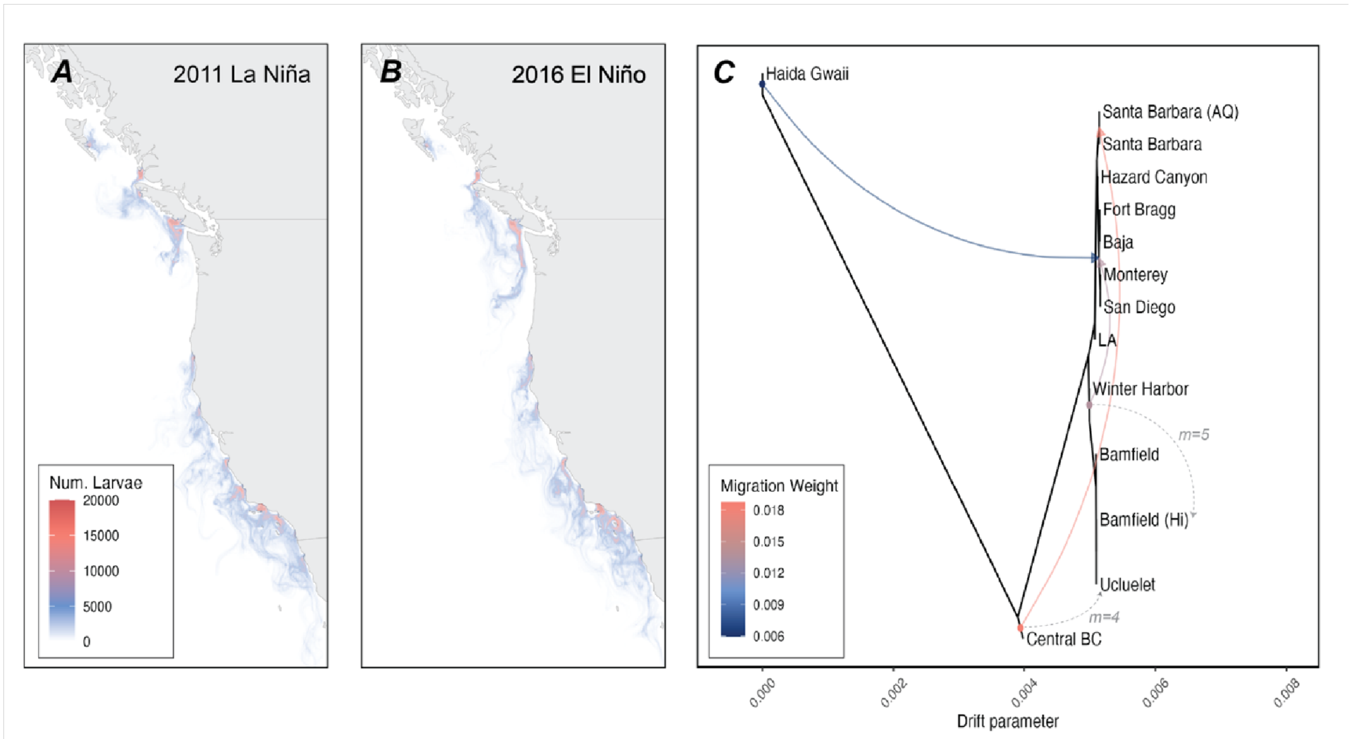


**Figure S14:** *Oceanographic Particle Simulation and TreeMix Analysis*
Representative results from a 10-week (summer) oceanographic particle simulation with Opendrift highlight decreased and increased southward dispersal in El Nino **(A)** and La Nina years **(B)**, respectively. Treemix analysis also highlights genetic signatures of north-south dispersal. These results are based on Treemix results using a consensus tree generated with 500 bootstrap replicates and 3 migration events. See **Supp. Figure 15** for bootstrap support (ranging from 74-100%). Manually drawn dashed lines indicate the additional migration edges inferred by Treemix when using 4 and 5 migration events.

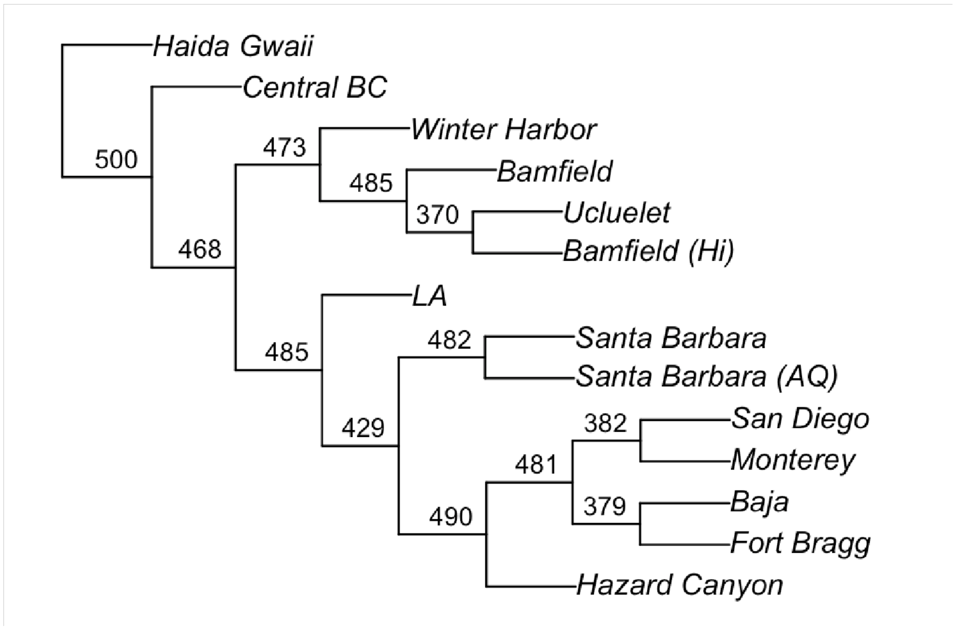


**Figure S15:** Phylogenetic tree generated with TreeMix showing bootstrap support from 500 replicate runs. This tree was used as a basis before inferring migration edges. To calculate these, we omitted the sample size correction as runs with this correction do not resolve any populations other than our most divergent samples in Haida Gwaii and Central BC. This highlights the similarity of populations south of the strong genetic split. However, the specific branching events in this tree should not be overinterpreted. We use this analysis to highlight a consistent pattern of north to south migration.
